# Potential of HLA-E-targeting diabodies to induce lysis of HIV-1-infected cells by CD8^+^ T cells

**DOI:** 10.64898/2026.04.28.721204

**Authors:** Srona Sengupta, Niklas Bachmann, Josephine Zhang, Jacqueline Douglass, Nathan L. Board, Fengting Wu, Milica Moskovljevic, Sarah DiNapoli, P. Aitana Azurmendi, Yasmine Tabdili, Madison Reed, Jeanna Yu, Emily Han-Chung Hsiue, Bailey Kim, Hao Zhang, Sandra B. Gabelli, Janet D. Siliciano, Robert F. Siliciano

**Author notes:** Address correspondence to: Robert Siliciano, Department of Medicine, Johns Hopkins University School of Medicine, 733 N. Broadway, Baltimore Maryland, 21205, USA;. These authors contributed equally to this work (Srona Sengupta, Niklas Bachmann). Discovery Chemistry, MRL, Merck & Co., Inc., Rahway, NJ 07065, USA.

## Abstract

A long-lived reservoir of cells harboring intact HIV-1 provirus persists throughout decades of antiretroviral therapy and can give rise to rapid viral rebound after treatment interruption. Some cure strategies employ cytotoxic T lymphocytes (CTL) to target this reservoir; however, the applicability and efficacy of immunotherapeutic strategies involving MHC class I-restricted CTL is limited by the polymorphic nature of MHC class I molecules and their downregulation by HIV-1 Nef. The non-polymorphic non-classical class I molecule HLA-E is stably expressed on HIV-1-infected CD4^+^ T cells and presents a potential universal target. We generated a single-chain diabody RLP-13 that redirects CTLs to target cells presenting a well-characterized peptide derived from *Mycobacterium tuberculosis* in the context of HLA-E. We verified the affinity and specificity of RLP-13. Through co-culture experiments, we confirmed that RLP-13 mediates polyfunctional, HLA-agnostic CTL responses. Using an HIV-1 reporter construct encoding the target peptide, we demonstrated robust and specific elimination of the HIV-1-expressing cell population. This proof-of-concept study shows that HLA-E antigens are promising immunotherapeutic targets that can bypass the limitations of classical MHC class I antigens – allelic variation and downregulation – and that such bispecific antibodies recognizing HIV-1-derived HLA-E binding epitopes could induce elimination of productively infected cells.

**Summary:** Sengupta, Bachmann et al. utilize a novel HLA-E-restricted CD3-engaging single-chain diabody to induce antigen-specific polyfunctional CTL-responses that are HLA-type-independent. They further show that such biologics have potential to eliminate HIV-1-infected cells by targeting HLA-E-binding epitopes encoded in the HIV-1 provirus.

## Introduction

T cell receptors (TCRs) recognize peptide fragments that are derived from pathogen proteins and bound to highly polymorphic major histocompatibility (MHC) molecules. Typically, peptides derived from viruses or intracellular bacteria are recognized by CD8^+^ T cells in the context of classical MHC class Ia molecules (HLA-A, B, C), whereas peptides derived from pathogen proteins present in the extracellular space and picked up by antigen-presenting cells are recognized by CD4^+^ T cells in the context of polymorphic MHC class II molecules (HLA-DR, DP, and DQ) (reviewed in (Blum et al., 2013; Neefjes et al., 2011)). In contrast to these highly polymorphic MHC molecules, HLA-E is a non-polymorphic, non-classical MHC class Ib molecule that shares homology with Qa-1 in mice and Mamu-E alleles in macaques (Sharpe et al., 2019). The two main alleles of HLA-E, HLA-E*01:01 and HLA-E*01:03, are present at similar population frequencies and differ only by the presence of either an arginine or glycine at position 107, which is located outside of the peptide binding groove (Strong et al., 2003) and thus not thought to influence the nature of peptides bound to HLA-E variants (Ottenhoff and Joosten, 2019). Similar to MHC-Ia, HLA-E is expressed on all nucleated cells, but at relatively low levels on the cell surface (Braud et al., 1998b).

The primary role of HLA-E is to prevent Natural Killer (NK)-cell mediated lysis of healthy, uninfected cells by presenting peptides derived from the leader sequence or signal peptide (SP) of HLA-A, B, C, and G molecules (Braud et al., 1998a; Braud et al., 1998b; van Hall et al., 2010). These peptide:HLA-E (p:HLA-E) complexes bind to members of the CD94/NKG2 family of receptors expressed on NK cells, with a particularly high affinity for the inhibitory NKG2A receptor (Ottenhoff and Joosten, 2019), resulting in a net inhibitory effect on NK cell-mediated lysis (Braud et al., 1998a). As some tumors and pathogens downregulate classical MHC class I molecules to limit recognition by CD8^+^ T cells, steady-state HLA-E-mediated presentation of signal peptides facilitates NK discrimination between normal “self” cells and “missing self” cells in which class I MHC molecules have been downregulated by a pathogenic process.

More recent research suggests that in addition to its canonical function in NK cell regulation, HLA-E and its rhesus and murine homologs can also present peptides derived from pathogens to CD8^+^ T cells. Implicated pathogens include *Mycobacterium tuberculosis* (*M. tb*) (Hansen et al., 2018; van Meijgaarden et al., 2015), along with several viruses: HIV-1 (Hannoun et al., 2018; Yang et al., 2021), EBV (García et al., 2002; Jørgensen et al., 2012), CMV (Mazzarino et al., 2005; Pietra et al., 2003), SARS-CoV-2 (Yang et al., 2023), and influenza (Hogan et al., 2023). Peptides bound to HLA-E have been eluted from *M. tb*-infected cells, and multiple HLA-E-associated *M. tb* peptides have been shown to stimulate CD8^+^ T cell responses (McMurtrey et al., 2017). Mamu-E restricted CD8^+^ T cell responses were also induced by vaccination using a novel rhesus cytomegalovirus vaccine vector (RhCMV68-1), which delivers recombinant simian immunodeficiency virus (SIV) genes. Remarkably, vaccination of macaques with the RhCMV68-1 vaccine and subsequent infection with the highly pathogenic SIVmac239 strain led to complete clearance of SIV infection in 55% of macaques (Hansen et al., 2011; Hansen et al., 2013a; Hansen et al., 2009). The primary immune correlate of vaccine-mediated protection was identified as an HLA-E-restricted, SIV-specific CD8^+^ CTL response (Hansen et al., 2013b; Hansen et al., 2016). These studies suggest that the diversity of the HLA-E-bound peptide repertoire post-infection is larger than previously appreciated and includes certain pathogen-derived peptides. Accordingly, HLA-E-directed strategies could be harnessed for the development of enhanced vaccine and/or immunotherapeutic strategies for various pathogens.

Using a highly diverse phage-display library, we previously identified antibody single-chain variable fragments (scFv) that recognize specific HIV-1 peptides bound to HLA-A:02*01. These HIV-1-specific scFv fragments could be incorporated into a single-chain diabody, a form of bispecific antibody in which one domain is specific for CD3ε, and the other domain binds to a pMHC-I complex. These TCR-mimic diabodies induced activation of CD8^+^ T cells and killing of HIV-1-infected cells (Sengupta et al., 2022). We hypothesized that this same strategy could be used to generate TCR-mimic bispecific antibodies specific for HIV-1 peptides bound to HLA-E for the purpose of redirecting CD8^+^ T cells of all specificities to kill HIV-1-infected cells bearing the relevant p:HLA-E. Given the nearly monomorphic nature of HLA-E compared to classical MHC-I alleles, such a bispecific reagent would have the ability to redirect CD8^+^ T cells from nearly all individuals to recognize and kill HIV-1-infected cells.

Two key challenges exist in constructing such bispecific agents. First, pathogen-derived peptides may not successfully compete with the natural ligands (signal peptides (SP) of MHC-I) for binding to HLA-E (Lin et al., 2023). Second, the number of pathogen-derived peptides that can bind with high-affinity to HLA-E is limited (Braud et al., 1997). Recently, however, pathogen-derived peptide-HLA-E complexes have been structurally characterized (Walters et al., 2018). One such complex involves the peptide Mtb44 (RLPAKAPLL), derived from the *inhA* gene of *M. tb*, which binds with high affinity to HLA-E (Walters et al., 2018). This epitope elicits CD8^+^ T cell responses in individuals with latent or active tuberculosis infections or in Bacillus Calmette-Guerin (BCG)-vaccinated children (Joosten et al., 2010).

The second characterized pathogen-derived pMHC complex involves the HIV-1 Gag epitope, HIV-Gag.275 (RMYSPTSIL), a NetMHC-predicted epitope that is homologous to the SIV Gag-derived SIV-Gag.276 (RMYNPTNIL) peptide, one of the two supertopes recognized by 100% of protected macaques in the RhCMV68-1 vaccine studies. The crystal structures of both of these peptides in complex with HLA-E*01:03 have been described (Walters et al., 2018). We reasoned that these particular high-affinity p:MHC could be used as ligands for phage display approaches and were able to successfully generate an HLA-E specific bispecific reagent. Here, we show that peptides known to bind with high affinity to HLA-E, when expressed in the context of HIV-1-infection, can be recognized by TCR-mimic bispecific antibodies leading to robust killing of HIV-1-infected cells.

## Results

### HLA-E is upregulated on HIV-1 infected cells

Redirected killing strategies involving HLA-E require that its surface expression on the target cell population is sufficient for CTL recognition. Therefore, we first examined changes in surface levels of HLA-E and classical class I molecules on primary CD4^+^ T cells following activation and infection with an HIV-1 reporter virus. We isolated CD4^+^ T cells from HIV-1-seronegative individuals, and either cultured them in RPMI-based media alone (no treatment condition) or activated them for 72 hours with anti-CD3/anti-CD28 beads in IL-2-supplemented media. Activated CD4^+^ T cells were then infected with an HIV-1 NL4-3 *Δenv*-eGFP reporter virus that expresses all viral genes except for the envelope (*env*) gene, which was truncated by an inserted eGFP cassette. The dual CCR5- and CXCR4-tropic *env* gene from the HIV-1 strain 89.6 was provided in trans during virus production. At 3 days post-infection, cells were sorted into GFP-negative and GFP-positive fractions in order to compare the effect of HIV-1 infection on surface levels of HLA proteins. Activation induced greater surface levels of HLA-A2, as can be seen by the significantly higher mean fluorescence intensity (MFI) of activated GFP-negative cells compared to control cells that had not been treated with anti-CD3/anti-CD28 beads (NT, not treated) (Figure 1A).

**Figure 1.**
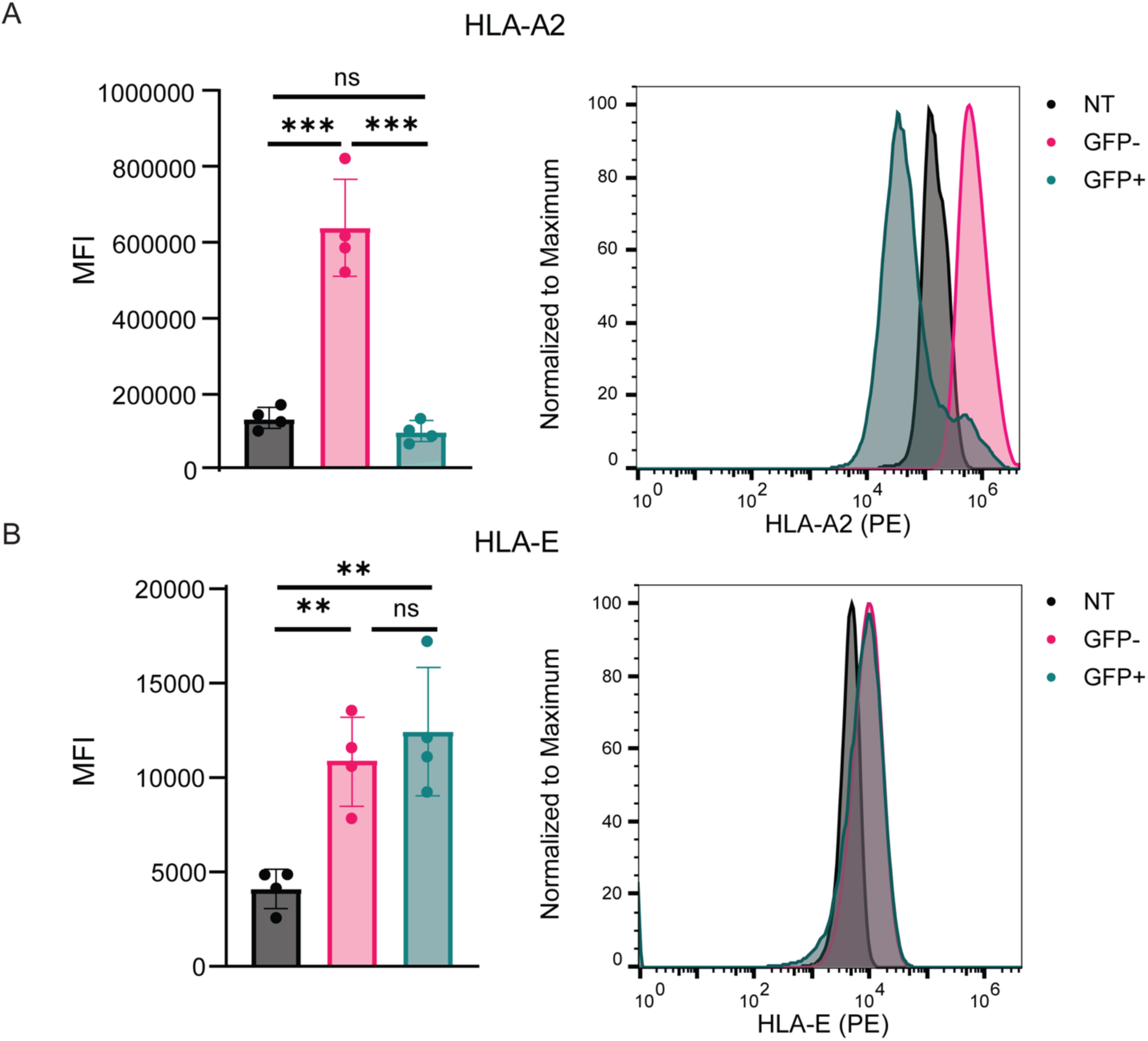
Surface MHC levels on uninfected or HIV-1-infected primary CD4^+^ T cells. Mean fluorescence intensity (MFI) of untreated (NT) or PHA-activated (GFP^-^, GFP^+^) primary human CD4^+^ T cells. Activated cells were infected with the HIV-1 reporter construct NL4-3 *Δenv*-eGFP and sorted into a GFP-negative and GFP-positive population. Cells were stained with lineage and viability markers and with PE-conjugated anti-human HLA-A2 (A) or PE-conjugated anti-human HLA-E (B), and the MFI of the viable CD3^+^CD8^-^ population was measured by flow cytometry. Each dot on the bar plot represents the average MFI for one HIV-negative donor (n = 4). Normalized histograms from one representative experiment are shown on the right. Statistical significance was determined through unpaired two-tailed t-test; ns: no significance (p-value > 0.05); **: p-value < 0.005; ***: p-value <0.0005.

However, in activated and productively infected cells (GFP^+^ population), the level of HLA-A2 was decreased by approximately 86% compared to the activated uninfected (GFP-) cells, indicating that HIV-1-infection induces downregulation of HLA-A2 (Figure 1A). This result is consistent with previous studies showing that the HIV-1 Nef protein downregulates MHC class I surface levels (Collins et al., 1998). For the non-classical HLA-E, surface levels were again higher in activated cells compared to cells that had not been activated, and no reduction was observed in the GFP^+^ population relative to GFP^-^ cells (Figure 1B). Thus, while abundance of the classical MHC class I molecule HLA-A2 is decreased on the surface of actively infected CD4^+^ T lymphoblasts, HLA-E levels remain stable upon infection.

### A phage display library can be used to isolate specific HIV-1 pHLA-E and M. tb pHLA-E scFvs

Given adequate levels of HLA-E on HIV-1-infected cells, we used a previously-generated phage display library with a diversity of ∼3.6 x 10^10^ unique clones (Miller et al., 2019; Skora et al., 2015) to screen for scFv-bearing phage that specifically bound HLA-E*01:03 in complex with the HIV-Gag.275, or SIV-Gag.276 peptides, or, as a control, the Mtb44 peptide (Figure 2A). Using the pan-MHC-I specific antibody W6/32 (Douglass et al., 2021), we confirmed correct folding of peptide-HLA-E (p:HLA-E) monomers, which we used to enrich for phage bearing scFv specific for the relevant p:HLA-E complexes through phage display panning. Our approach was similar to that taken previously (Douglass et al., 2021; Hsiue et al., 2021; Sengupta et al., 2022): negative selection with streptavidin beads bound to biotinylated p:HLA-E complexes bearing irrelevant peptides and cells bearing non-target epitopes bound to HLA-E, and positive selection with decreasing amounts of cognate p:HLA-E complexes. Five rounds of panning yielded 109 unique scFv clones for HIV-Gag.275, 79 unique clones for SIV-Gag.276, and 109 unique clones for Mtb44. However, when individual phage clones were tested for binding to respective biotinylated cognate pHLA-E bound to streptavidin-coated plates, minimal binding was observed, suggesting overly stringent selection in later rounds of panning or outgrowth of phage clones with growth advantage.

**Figure 2.**
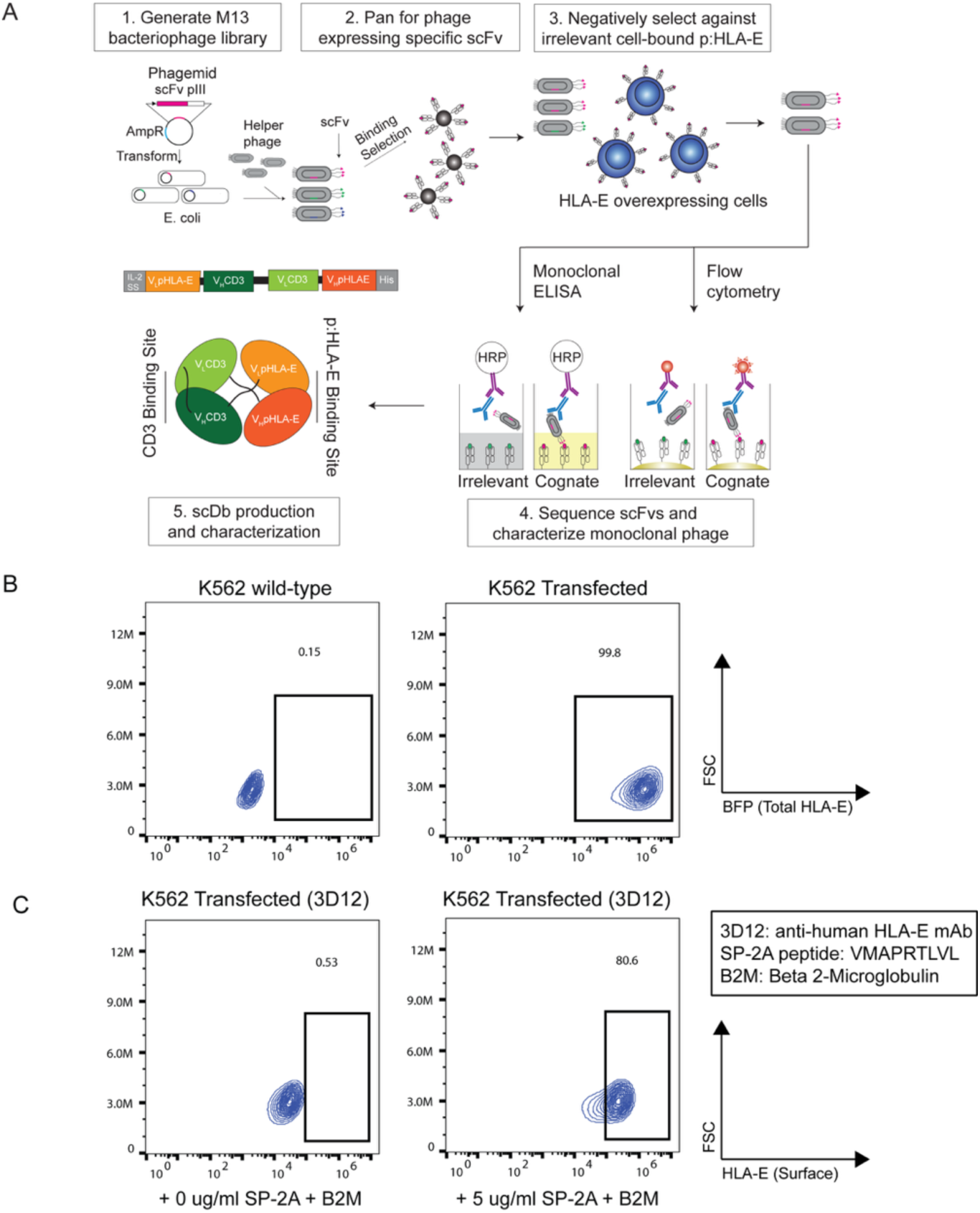
Overview of phage panning and design of single-chain diabody. (A) An M13 bacteriophage library expressing 3.6x10^10^ single-chain variable Fragments (scFvs) was generated and screened as described previously (Miller et al, J Biol Chem 2019). Negative selection to remove clones binding irrelevant p:HLA-E complexes was performed by phage panning on HLA-E-overexpressing cells pulsed with irrelevant peptides. Positive selection for clones binding the relevant p:HLA-E complex was performed by panning with microbeads coated with cognate peptide:MHC. After up to 5 rounds of panning, individual clones were assessed for specificity through monoclonal phage ELISA and flow cytometry, and specific clones were sequenced. Heavy and light chain sequences of candidate scFvs were inserted into a CD3ε-engaging scDb constructs flanked by an IL-2 secretion signal and a 6xHis-tag. (B) Characterization of HLA-E-overexpressing K562 cells used in negative selection. Flow cytometry contour plots show that relative to control wild type K562 cells, there was uniformly high BFP expression (a marker for total HLA-E levels) in K562 cells that were transfected with an HLA-E overexpression construct. (C) Surface HLA-E levels on HLA-E-overexpressing clone 3.3 of the transfected K562 cells shown in (B). Cells were left unpulsed (left) or pulsed with 5 μg/mL the HLA-A2 signal peptide SP-2A (VMAPRTLVL) peptide and 10 μg/mL beta 2-microglobulin for 4 hours, and stained for flow cytometry with PE-labeled anti-human HLA-E antibody (clone 3D12).

We therefore modified our panning protocol. In our second attempt at panning for p:HLA-E targets, we included an HLA-E overexpressing cell line (Figure S1A) for negative selection. As HLA-E is expressed on the cell surface of resting and active CD4^+^ T cells at lower levels compared to classical MHC-Ia molecules (Figure 1), we reasoned that cells typically used for negative selection (Jurkat T cells or primary CD4^+^ T cells) would not likely display sufficient p-HLA-E complexes for adequate negative selection. Therefore, we generated an HLA-E overexpressing line from K562 cells. The K562 cell line is a multipotential leukemic cell line derived from a patient with chronic myelogenous leukemia(Boegel et al., 2014; Lozzio and Lozzio, 1979; Sutherland et al., 1985). These cells lack expression of MHC I molecules. To generate a cell line that constitutively expresses high levels of HLA-E, we transfected K562 cells with a construct containing the HLA-E*01:03 coding sequence linked at the C-terminus to a BFP tag, with the sequences separated via a T2A sequence which induces ribosomal skipping. To allow constitutive expression, these sequences were placed downstream of a CMV promoter. In the resulting cell line, total intracellular and extracellular HLA-E expression could be assessed via the BFP tag (Figure 2B). This cell line, termed K562-3.3, could be pulsed with the signal peptide from HLA-A2 (SP-2A, or VMAPRTLVL), which stabilized surface HLA-E expression (Figure 2C). A similar effect was observed when pulsing these cells with any of the target peptides, although to a lesser degree with HIV.Gag275 and SIV.Gag276 (Figure S2). During the panning process, we pulsed K562-3.3 cells with saturating levels of the SPs representing the physiologic ligands of HLA-E, SP-1A (VMAPRTLLL) and SP-2A (VMAPRTLVL), as well as peptide mimics with single amino acid substitutions in the target peptides, to enable more stringent negative selection against scFv capable of recognizing HLA-E complexed with endogenous ligands. In addition to the inclusion of this cell line, a second key difference during HLA-E phage panning was positively selecting phage using 10-fold lower levels of cognate pMHC bound to streptavidin beads, with the goal of identifying higher-affinity scFv (Table S1). Using this modified panning protocol, we were able to identify 187 clones for HIV-Gag.275, 81 clones for SIV-Gag.275, and 192 clones for Mtb44 from rounds 3 and 4 of panning. Several scFvs appeared to bind to cognate HIV-Gag.275, SIV-Gag.276, and Mtb44 at levels >1.5 times the binding to the non-cognate SP-2A control. Particularly for Mtb44, some isolates exhibited more than 2-fold increased binding to the cognate target (Figure S3C).

### Pathogen-derived pHLA-E scFvs can be converted to single-chain diabodies that induce specific T cell activation

The scFv sequences of phage “hits” were converted into a single-chain diabody (scDb) format (Douglass et al., 2021; Hsiue et al., 2021) consisting of a single 6xHis-tagged polypeptide chain containing the heavy and light chain variable regions of the two scFv fragments separated by flexible glycine repeat linkers (Figure 2A). In these constructs, the scFv of the anti-pMHC domain derived from phage panning flanks the scFv of the anti-CD3ε clone UCHT-1 (Douglass et al., 2021). Binding of the scDb to CD3 on the effector cell and the HIV-1 pMHC on the infected target cell should tether the effector to the target to form an immune cytolytic synapse, with activation of the effector cell as assessed by CD69 and CD107 upregulation, release of cytokines and chemokines such as IFNγ and MIP1β (Douglass et al., 2021; Hsiue et al., 2021), and target cell lysis.

To test the ability of these scDbs to bind to the target pHLA-E complexes, we incubated the scDbs with biotinylated pHLA-E immobilized to streptavidin plates and probed for the binding of the scDb using anti-His secondary antibodies (see Methods). We did not observe binding of any of the HIV.Gag275 scDbs tested (n = 50), but out of n = 27 SIV.Gag276-targeting constructs tested, we did observe binding by the constructs RMY276-3, RMY276-61, and RMY276-74 to the target SIV.Gag276 pMHC by ELISA (Figure 3A). Of note, we also observed binding of these scDbs to HIV.Gag275. This was expected as the sequence RMYSPTSIL (HIV) is homologous to the SIV sequence RMYNPTNIL, and the most solvent exposed residue (P6) is the shared Threonine on both pMHC complexes (Walters et al., 2018). We tested the functionality of these SIV-specific scDbs by co-culturing CD8^+^ T cells from HIV-negative donors with K562-3.3 cells pulsed with 10 µM of the SIV.Gag276 peptide, HIV.Gag275, SP-2A peptide or no peptide (negative control). We assessed effector cell activation by cell-surface expression of the cytolytic degranulation marker CD107 (Seder et al., 2008) (Figure 3B). However, the most promising SIV.Gag276-targeting scDb, RMY276-61 induced upregulation of CD107 in each co-culture condition regardless of the target peptide (Figure 3B), potentially indicating non-specific binding to HLA-E.

**Figure 3.**
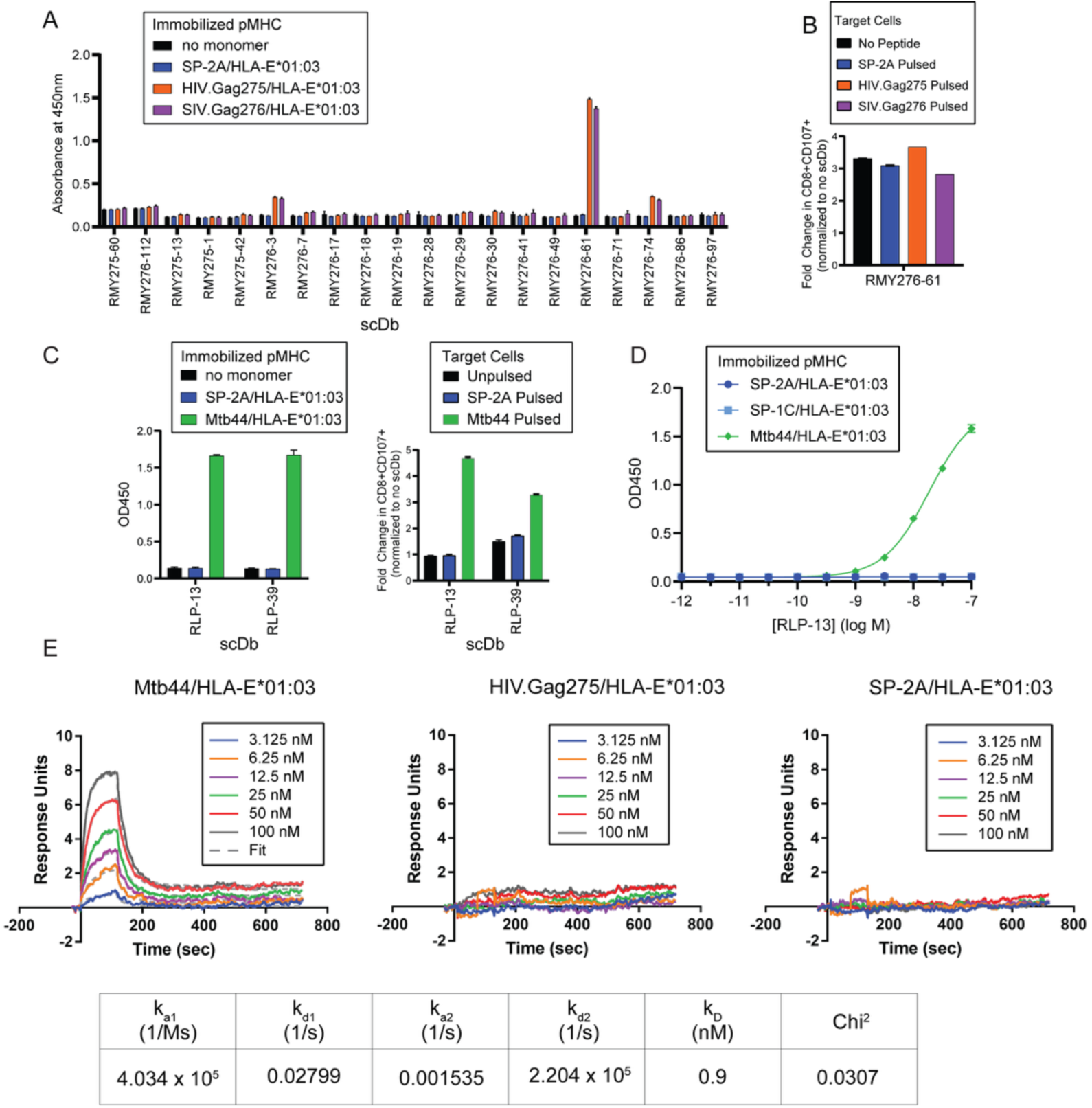
Specificity and affinity of individual scDb clones. (A) Absorbance from scDb binding ELISA using immobilized cognate and non-cognate peptide:HLA-E mononmers. (B) CTL-stimulation of scDb construct RMY276-61 in co-cultures with K562-3.3 cells pulsed with indicated peptides, as measured by surface levels of CD107. (C) Specificity of scDb constructs RLP-13 and RLP39 measured in peptide:MHC-binding ELISAs (left) and of corresponding scDbs in co-culture assays with peptide-pulsed K562-3.3 cells (right). (D) Binding of RLP-13 against immobilized HLA-E monomers loaded with SP-2A (VMAPRTLVL), SP-1C (VMAPRTLLL) or Mtb44 (RLPAKAPLL) measured by ELISA. (E) RLP-13 binding to RLP was measured using SPR with multicycle kinetics at concentrations ranging from 3.125 nM up to 100 nM. The two-state reaction model fitting is shown in dashed grey lines. No binding was measured against SP-2A and HIV.Gag275 for all concentrations tested. Calculated binding kinetics coefficients are given below.

Although screening of an extensive phage library did not yield functionally active scDbs specific for the HIV-1 or SIV Gag peptides, we did find two scDbs specific for the *M. tb*-derived peptide Mtb44 (RLPAKAPLL). The corresponding scDbs, RLP-13 and RLP39, both exhibited specific binding to the immobilized cognate pHLA-E and not to the no-monomer or irrelevant monomer (SP-2A/HLA-E*01:03) control (Figure 3C). The RLP-13 scDb induced greater and more specific CTL activation, as measured by surface levels of CD107, compared to the RLP39 scDb (Figure 3C). Additional testing of RLP-13 via a titration ELISA highlighted its specificity. RLP-13 scDb only bound to immobilized Mtb44/HLA-E*01:03 and not to the irrelevant monomers SP-2A/HLA-E*01:03 or SP-1C/HLA-E*01:03 (VMAPRTLIL) even at high concentrations of the RLP-13 scDb (Figure 3D). The specificity of RLP-13 was additionally interrogated using positional scanning variant peptides, in which each residue of the Mtb44 peptide was substituted with 19 other possible amino acids. K562-3.3 cells were pulsed with the resulting library of 171 variant peptides. To assess the impact of single residue substitutions on the stability of the peptide-MHC interaction, we performed HLA-E surface staining of cells pulsed with each variant. In accordance with a previously published study (Ruibal et al., 2020), residues in position 2, 7 and 9 were the least permissive to amino acid substitutions, as evident from reduced levels of HLA-E on the cell surface (Figure S4A). Pulsed cells were also co-cultured with RLP-13 and CD8^+^ T cells, and secretion of MIP-1β into the supernatant was assessed to determine which amino acid changes affect recognition of the pMHC by RLP-13. In addition to those residues that are crucial for the peptide binding to HLA-E, RLP-13-induced CTL activation was significantly decreased in co-cultures with K562-3.3 cells pulsed with peptide variants containing substitutions in positions 1, 4 and 6, suggesting that these residues were critical for the interaction between RLP-13 and the peptide:HLA-E complex. Positions 3, 5 and 8 tolerated more substitutions (Figure S4B).

Due to robust activity and specificity of RLP-13, we pursued additional functional characterization of this scDb. Surface plasmon resonance (SPR) analysis demonstrated RLP-13 binding to Mtb44/HLA-E*01:03 in a two-state reaction model with a calculated equilibrium constant (K_d_) of 0.9 nM (Figure 3E). This reflects an affinity comparable to and in some cases higher than the affinity of previously reported scDbs (Douglass et al., 2021; Hsiue et al., 2021; Sengupta et al., 2022). The first association rate constant (k_on_) was 4.034 x 105 M^-1^s^-1^ and a dissociation rate constant (k_off_) of 27.99 x 10^-3^ M^-1^s^-1^. The second association rate constant (k_on_) was 0.001535 M^-1^s^-1^, and the dissociation rate constant (k_off_) was 2.2 x 10^-5^ M^-1^s^-1^. The fit with a two-state reaction model suggests a conformational change that stabilizes the complex. Other scFvs, such as V2 against KRAS G12V bound to HLA-A3, also display this behavior (Douglass et al., 2021). RLP-13 did not bind to either HIV-Gag.275/HLA-E*01:03 or SP-2A/HLA-E*01:03 (Figure 3E). Overall, this analysis underscores the high affinity of RLP-13 to the relevant *M. tb* pMHC, which would be required for specific CTL activation in the setting of much more abundant non-cognate pMHC.

### RLP-13 induces robust CTL killing of cognate pHLA-E bearing cells with high sensitivity

To further assess the specificity and functional activity of RLP-13 in vitro, we performed co-culture killing assays using primary CD8^+^ T cells and K562 cell lines that were transfected to stably overexpress covalently linked cognate peptide (Mtb44/HLA-E*01:03) or non-cognate peptide (SP-2A/HLA-E*01:03) pMHC (Walters et al., 2018) (Figure 4, A and B). We observed specific CTL activation and dose-dependent lysis of up to ∼92% cognate target cells at the highest concentration of RLP-13 tested (3nM), with no significant loss in viability of non-cognate target cells (Figure 4C). H2-mu, an irrelevant control scDb specific for a mutant p53^R175H^ epitope in the context of HLA-A*02:01 (Hsiue et al., 2021), did not cause any reduction in viability in the cognate or non-cognate target cell population. In addition to canonical markers of CTL degranulation (granzyme A, Perforin, sFasL) (Figure 4D), RLP-13, but not irrelevant control scDb, induced the release of cytokines TNF-α, INFγ, IL-2, IL-4, IL-6, IL-10, IL-17A, chemokine MIP-1β, as well as Granzyme B and sFas in a dose-dependent manner (Figure S5).

**Figure 4.**
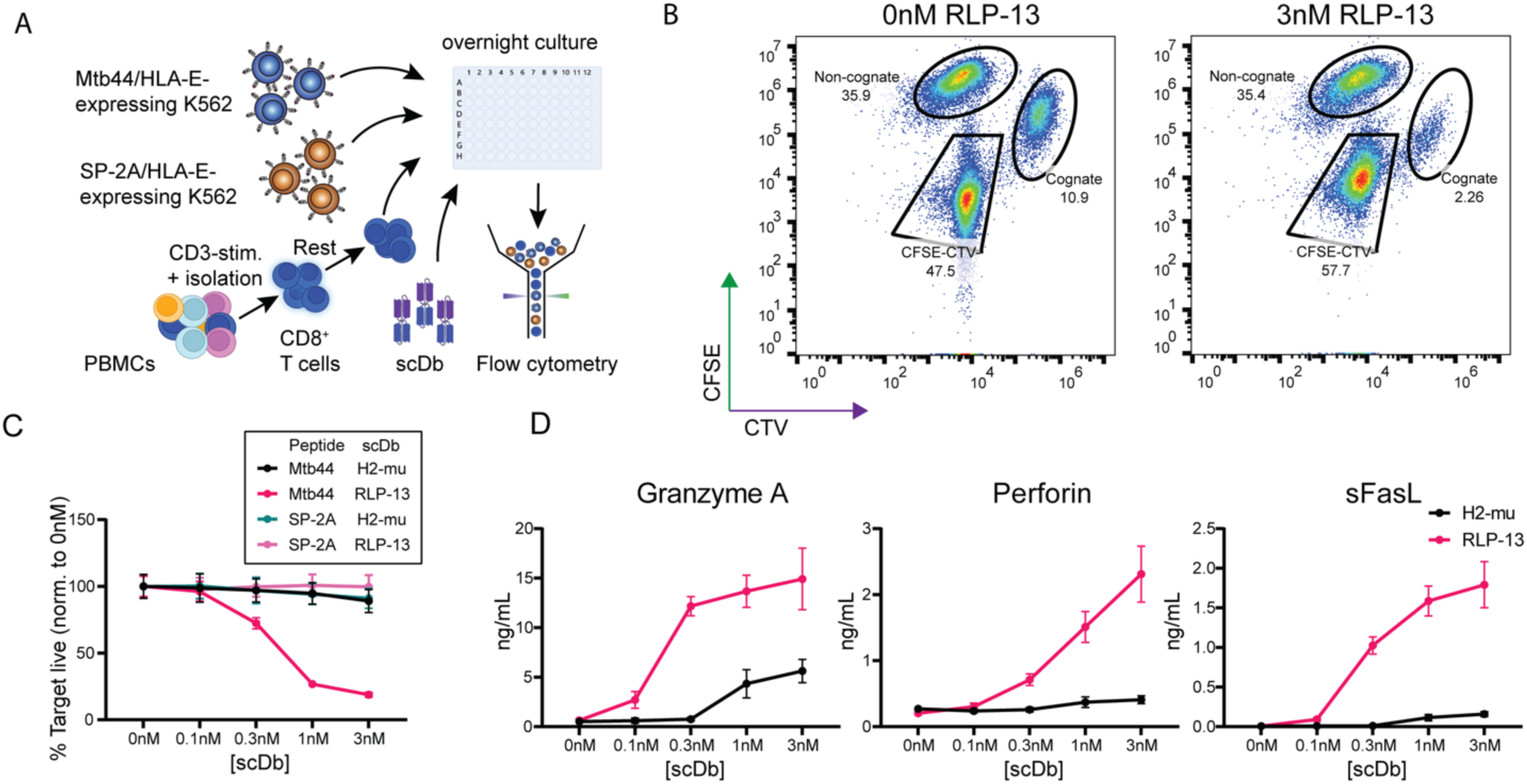
Specificity of RLP-13 in dual-color co-cultures. (A) Setup of dual-color co-cultures: K562 cells transfected with covalently linked peptide:HLA-E expression constructs, presenting either cognate (Mtb44) or non-cognate (SP-2A) peptide, are stained with CellTrace Violet (CTV) or CFSE, respectively, and cultured for 18 hours with pre-expanded CD8^+^ T cells and different concentrations of RLP-13 at a ratio of 4:1:3 (Effector : Cognate target : Non-cognate target). Cells are then stained with lineage and activation markers and viability dye for analysis by flow cytometry. (B) Representative flow plot of viable single cell populations from co-culture wells without (left) or with (right) RLP-13. Indicated cognate and non-cognate gated population frequencies were used to calculate the relative viability of target cell populations. (C) Target cell viability of indicated cognate and non-cognate populations from co-cultures with various concentrations of either RLP-13 (red and pink lines) or irrelevant control scDb H2-mu (black and green lines). Viability was calculated by normalizing the frequency of the indicated target cell populations to that observed in co-culture wells without any scDb added. Assay was repeated five independent times across two HIV-negative donors. Shown measurements are averaged over eight technical replicates from one representative experiment. (D) Concentrations of the indicated effector molecules in the supernatants of co-cultures shown in (C) measured using the LegendPlex CD8/NK cell cytokine panel kit.

Effective TCR mimic reagents must detect pathogen-derived peptide-HLA-E complexes presented at low copy number on target cells. Therefore, we performed peptide titration experiments. The target cells in these co-cultures were peripheral blood mononuclear cells (PBMCs) from HIV-negative donors who were homozygous for either HLA-E*01:01 or HLA-E*01:03. PBMCs were pulsed with the Mtb44 peptide at concentrations ranging from 10 pM to 10 µM, and cultured with pre-expanded autologous CD8^+^ T cells and with or without RLP-13 (Figure 5A). The RLP-13 scDb induced specific, polyfunctional T cell responses against PBMCs pulsed with picomolar concentrations of the relevant peptide for both HLA-E alleles (Figure 5B). The levels of CTL effector molecules detected in the corresponding supernatants were comparable between co-cultures using donor cells homozygous for either HLA-E allele, supporting the hypothesis that antigen recognition by RLP-13 occurs at similar efficiencies regardless of the HLA-E allele (Figure 5, C and D).

**Figure 5.**
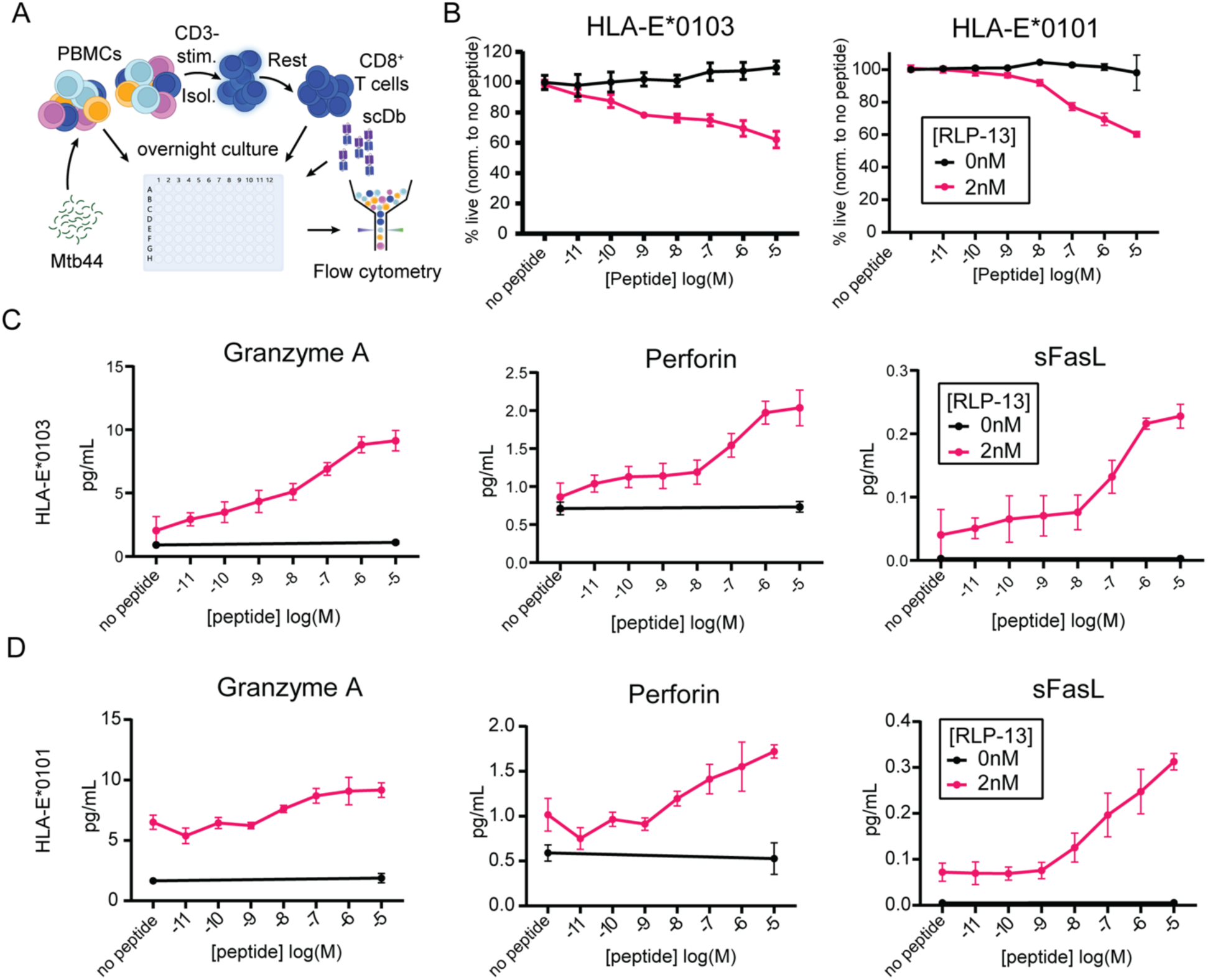
Efficacy and sensitivity of RLP-13 in primary cell co-cultures for different HLA-E alleles. (A) Overview of co-culture setup: PBMCs isolated from HIV-negative donors homozygous for either HLA-E*01:01 or HLA-E*01:03 were pulsed with various concentrations of Mtb44-peptide, and cultured with pre-expanded autologous CD8^+^ T cells and either 0 nM or 2 nM RLP-13 overnight at an effector-to-target ratio of 3:1. Cells were then stained with lineage and activation markers and viability dye. (B) Viability curves of target cells from co-cultures described in (A). Assay was performed four times across two donors. Shown measurements are averaged over at least six technical replicates from representative assays for each donor. (C) and (D) show effector molecule concentrations in supernatants from co-cultures in (B) measured using the LegendPlex CD8/NK cell cytokine panel kit.

### Killing of HIV infected cells bearing the appropriate epitope

The RLP-13 scDb showed high specificity and efficacy in redirecting CTLs to lyse HLA-E expressing target cells pulsed with Mtb44. However, peptide pulsed target cells may display the relevant p:MHC complexes at much higher levels than are typically present on the surface of infected cells (Sengupta et al., 2022). Therefore, we tested whether the scDbs could be used to eliminate primary CD4^+^ T cells productively infected with a recombinant HIV-1 reporter virus carrying the Mtb44 sequence. The nucleotide sequence encoding the Mtb44 peptide was inserted after the eGFP open reading frame of HIV NL4-3 *Δenv*-eGFP, separated by a T2A sequence to induce ribosomal skipping during translation and avoid interference of the Mtb44 peptide with eGFP function (Figure 6A). This likely results in delivery of the peptide into the cytoplasm and subsequent uptake into the ER. We then used this viral reporter construct (ΔEnv-Mtb44), as well as the original construct lacking the Mtb44-sequence (ΔEnv), in co-culture cytotoxicity assays to assess the specificity and efficacy of RLP-13-induced killing (Figure 6B).

**Figure 6.**
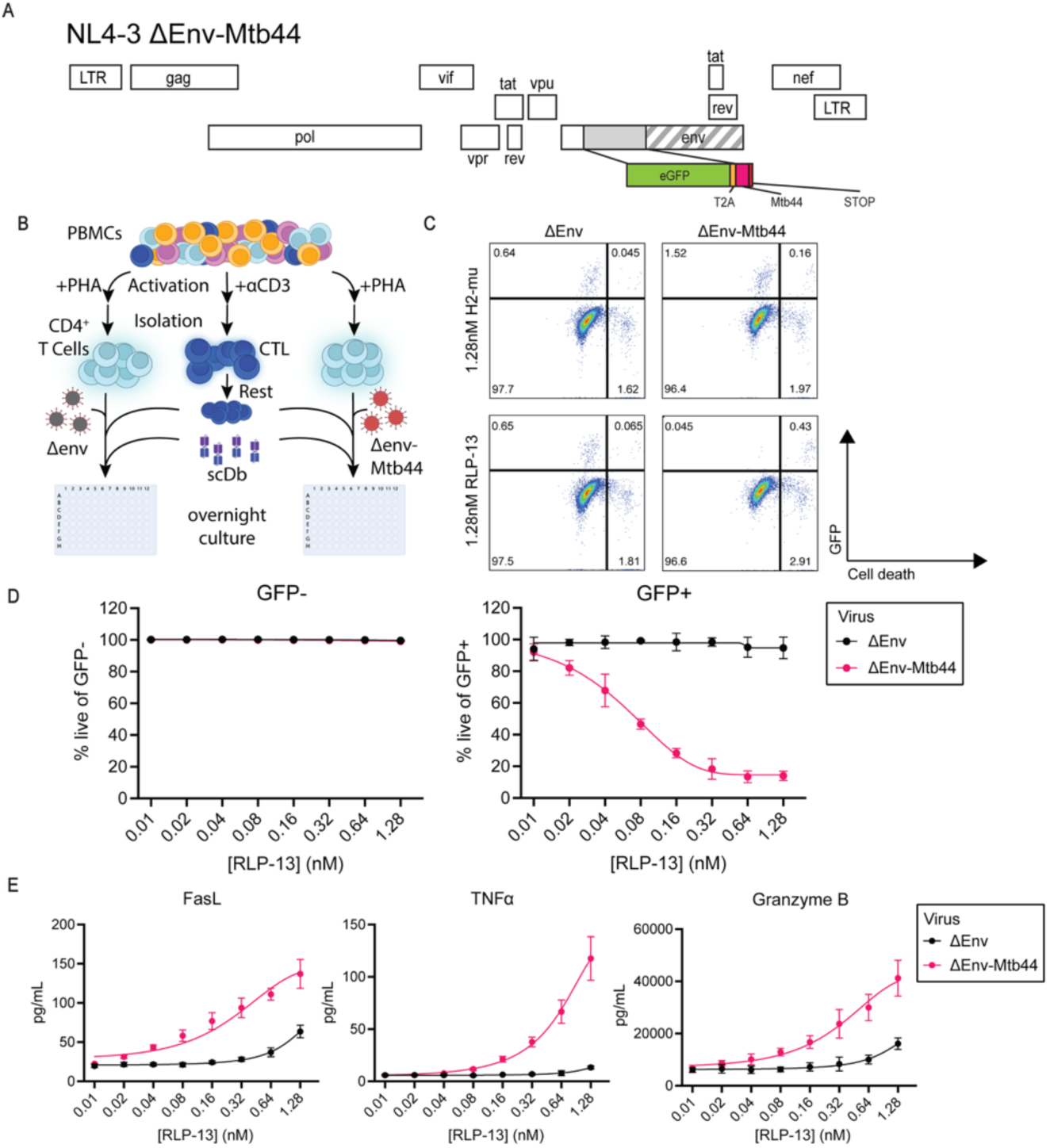
RLP-13 induces antigen-dependent lysis of cells expressing HIV-1 construct encoding for Mtb44 peptide (A) Outline of the NL4-3 *Δenv*-Mtb44 reporter construct. The provirus encodes for all HIV-1 gene products except *env*, which is truncated by an inserted eGFP reporter-cassette followed by a non-coding T2A-sequence and the DNA sequence encoding for the Mtb44-peptide. (B) Experiment setup: PBMCs from isolated from three HIV-negative donors. PBMCs were treated with PHA for 72 hours before infection with the indicated HIV-1 reporter construct without (ΔEnv) or with (ΔEnv-Mtb44) the insert encoding for the *Mtb44*-peptide. Effector cells were prepared by culturing autologous PBMCs with anti-human CD3 monoclonal antibody, and CD8^+^ T cells were isolated and rested for 7 days. Target and effector cells were co-cultured overnight at a ratio of 1:1 with varying concentrations of RLP-13. Cells were then stained for viability, lineage and activation markers and analyzed by flow cytometry. (C) Selected flow cytometry plots from a representative experiment as described in (B). Target cells were identified as CD3^+^CD8^-^ single cells. Each plot shows data from one well containing target cells infected with the indicated virus for each column and cultured with autologous CD8^+^ T cells and the indicated scDb for each row. (D) Frequency of viable GFP-positive (left) or GFP-negative target cells (right). PHA-treated, activated CD4^+^ T cells were infected with either NL4-3 *Δenv*-eGFP or NL4-3 *Δenv*-Mtb44 and cultured with autologous, pre-expanded CD8^+^ T cells and indicated concentrations of RLP-13. Target cells were identified as CD3^+^CD8^-^ single cells. GFP-expressing indicates active expression of integrated provirus. Viability was normalized to an irrelevant scDb (H2-mu) control. Measurements are averaged over four technical replicates. Two independent repeat experiments were performed for three HIV-negative donors; data shown is from one representative experiment. (E) Concentrations of the indicated effector molecules in the supernatant of co-cultures from (D).

Primary CD4^+^ T cells from HIV-negative donors were activated and infected with either of the two HIV-1 reporter constructs and co-cultured with autologous CD8^+^ T cells and varying concentrations of RLP-13 or irrelevant scDb overnight, and viability of GFP-positive and GFP-negative cell populations from the same wells were compared (Figure 6C). The GFP-expressing target cell population was reduced in an RLP-13 dose-dependent manner only in cells infected with the Mtb44-encoding construct, whereas no killing was observed in GFP-negative cells or cells infected with virus that did not contain the Mtb44-sequence (Figure 6, C and D). RLP-13 induced elimination of cognate target cells at RLP-13 concentrations as low as 10pM, with a maximum reduction of the GFP^+^ population of approximately 90% at 1.28 nM (Figure 6D). Co-culture supernatants were tested for a panel of CTL effector molecules including Granzyme B, TNFα and sFasL. In the presence of cells expressing the Mtb44-encoding construct, supernatant concentrations of these molecules increased in a robust and dose-dependent manner. In the absence of the Mtb44-peptide, changes in concentrations of CTL effector molecules secreted into the culture supernatant were only notable at the highest doses of RLP-13 tested (Figure 6E), although even in those conditions, no off-target killing was observed.

Together, these results suggest that HLA-E-binding peptides encoded by an HIV-1 provirus can be processed and presented in the context of HLA-E on the cell surface at levels sufficient for recognition by a specific scDb.

## Discussion

A major obstacle to curing HIV-1 infection is the persistent, long-lived population of cells harboring an intact HIV-1 provirus which can give rise to viral rebound within weeks of cessation of antiretroviral therapy. Previous studies have shown that the natural decay of this reservoir over decades of ART suppression of viremia is not sufficient to eliminate this reservoir (Crooks et al., 2015; McMyn et al., 2023; Siliciano et al., 2003); thus, additional intervention is needed to specifically target and eliminate HIV-1-infected cells. One potential method that is employed in the field of immune-oncology is the redirecting of cytotoxic T lymphocytes using bispecific reagents (Merz et al., 2024) (reviewed in (Shui et al., 2025) and (Nordstrom et al., 2022)). Here we describe the generation of a novel bispecific T cell engaging antibody construct specific for an HLA-E-bound *M. tb*-derived epitope. This reagent induced robust, highly specific killing of primary CD4^+^ T cells pulsed with the relevant peptide and of HIV-1-infected cells expressing the target peptide from an integrated provirus.

The potential of bispecific T cell engagers such as single-chain diabodies to induce cytolytic killing of cancer cells (Douglass et al., 2021; Hsiue et al., 2021; Lutterbuese et al., 2010; Löffler et al., 2000) or infected cells (Quitt et al., 2021; Sengupta et al., 2022) has been demonstrated in vivo and in vitro. By virtue of engaging the conserved CD3 component of the TCR receptor, these bispecific antibodies have the advantage of being able to redirect all T cells, regardless of TCR specificity, to target cells bearing the relevant epitope. Additionally, these engagers provide a quick, cost-efficient and controllable option for immunotherapeutic intervention that does not depend on personalized and lengthy processes as is the case for adoptive cell transfer approaches. Current bispecific antibody constructs are primarily used in the context of cancer immunotherapy, with a few more recent ongoing trials to target viral infections (reviewed in (Nyakatura et al., 2017; Shui et al., 2025)). Blinatumomab, teclistamab and tebentafusp-tebn are three examples of bispecific T cell engagers, including scDbs, that were approved by the US Food and Drug Administration for treatment of acute lymphoblastic leukemia, multiple myeloma or uveal melanoma, respectively. These bispecific antibodies typically target tumor-associated surface proteins such as CD19. In cases where tumor- or pathogen-associated surface proteins are not available for targeting, tumor-/pathogen-specific peptides presented in the context of MHC could be targeted through either scFvs or TCR constructs. Such MHC-restricted bispecific antibodies have been developed for both neoantigen peptides (*GUARDIAN-101 trial* for CLSP-1025, which targets p53^R175H^/HLA-A*02:01; ClinicalTrials.gov ID NCT06778863) and pathogen-derived peptides (*STRIVE trial* for IMC-M113V, which targets HIV Gag_77-85_/HLA-A*02:01; EudraCT number 2021-002008-11); these constructs are being tested in ongoing clinical trials.

While the repertoire of MHC-bound peptides offers additional potential immunotherapeutic targets, applicability of these treatments is limited by the polymorphic nature of MHC. Over 7,000 alleles for HLA-A have been reported to date, and over 17,000 alleles across all MHC class I and class II genes ((EBI), 2017a; (EBI), 2017b). These polymorphisms affect both the topology of MHC molecules, and thus the surface recognized by bispecific antibodies, but also the shape of the peptide-binding grove and consequently the repertoire of peptides that can be stably presented by each HLA molecule. Therefore, application of a specific scFv is typically restricted to a subset of people who express the exact HLA-allele that the bispecific antibody was designed for. HLA-A*02:01 is the most prevalent HLA allele, found in over 20% of the European-descendant population and over 10% in African and African-American populations (Olivier et al., 2023; Sanchez-Mazas et al., 2024). This allele is the target for several bispecific MHC-restricted therapeutics such as CLSP-1025 and IMC-M113V. However, a majority of people would not benefit from such treatments. The non-classical MHC class I molecule HLA-E is non-polymorphic, and the only amino acid difference between the two equally common alleles HLA-E*01:01 and HLA-E*01:03 is located outside of the peptide binding grove. Thus, all HLA-E molecules likely bind the same peptide repertoire at comparable affinities, offering an opportunity for universal application provided that appropriate target peptides can be presented by HLA-E.

Given the universal targeting enabled by HLA-E, and more recent evidence showing a role for HLA-E in presenting pathogen-derived peptides (Walters et al., 2018; Yang et al., 2021), we sought to generate a bispecific engager that could bind HLA-E on the surface of an HIV-infected cell and induce specific killing. Building on our prior work (Sengupta et al., 2022), and in contrast to prior studies that have developed therapeutics targeting structural surface antigens or linearized peptides presented by classical MHC class I molecules, we employed scFv phage panning to design a novel scDb specific to peptides bound to the non-classical MHC class I molecule HLA-E. We generated a new bispecific construct, RLP-13, that recognizes the high-affinity *M. tb*-derived Mtb44-peptide and validated its affinity and specificity through both biophysical and functional assays. We further showed that RLP-13 induces polyfunctional CTL responses and lysis of target cells homozygous for either of the two common HLA-E-alleles, HLA-E*01:01 and HLA-E*01:03, at similar levels and with similar sensitivity. These results underscore the universal targeting of these reagents regardless of HLA type. Finally, our findings from co-culture assays with primary cells expressing Mtb44-encoding HIV-1 reporter constructs indicate that endogenous or provirus-derived HLA-E-binding peptides are expressed and presented on the cell surface at levels sufficient for scDb-mediated CTL killing. Overall, this study shows that HLA-E presents a feasible immunotherapeutic target that allows for the design of reagents that are universally applicable, instead of being restricted to a small subpopulation of carriers of specific HLA-A/B/C alleles, and that allow for specific targeting of primary cells that endogenously express an appropriate HLA-E-binding peptide.

HLA-E based bispecific reagents not only address the allele-specific limitation of MHC-I targeting bispecific antibodies, but also MHC-I downregulation induced by some tumors and infections such as HIV-1 (reviewed in (Cornel et al., 2020; Kallingal et al., 2023)). This immune evasion mechanism can prevent recognition by CTLs, but also reduces inhibitory signals to NK cells, which may be activated when encountering cells lacking MHC class I (Ljunggren and Kärre, 1990). Surface expression of HLA-E, which binds to the inhibitory NK cell receptor CD94/NKG2A (Lee et al., 1998), can suppress NK cell activation and is therefore beneficial for the survival of the expressing cells, making HLA-E a promising, stably expressed target for bispecific reagents and other therapeutics. It is worth nothing that HLA-E surface levels of infected cells may vary between HIV-1 isolates (van Stigt Thans et al., 2019). However, our findings from co-culture experiments with HIV-1-infected cells indicate that even in the setting of HIV-1 Nef-induced MHC class I downregulation, RLP-13 could induce efficient elimination of the target cell population. Furthermore, this is evidence that the lower surface levels of HLA-E compared to classical MHC class I molecules are sufficient for robust recognition and scDb-mediated lysis.

Overall, this study provides a proof of concept that HLA-E-bound peptides can be utilized for immunotherapeutic targeting. Employing scFv phage panning, we generated RLP-13, a novel, highly specific scDb recognizing a pathogen-derived peptide in the context of both HLA-E*01:01 and HLA-E*01:03, indicating that such HLA-E antigens allow for universally efficient therapeutics design without being limited to only the population expressing a particular HLA allele. We demonstrated that RLP-13 induces an antigen-specific polyfunctional CTL response in cultures with primary cells pulsed with peptide concentrations as low as 10 pM. Our findings showed that non-specific CTL activation in the absence of cognate antigen only occurred at the highest tested concentrations of RLP-13 or the control scDb H2-mu, and no significant off-target toxicity was observed even at saturating scDb concentrations. Furthermore, we demonstrated the potential of HLA-E targeting in the context of HIV-1 infection. RLP-13 induced specific and robust target cell lysis of cells expressing the target peptide from the HIV-1 provirus, and was not affected by HIV-1 Nef-induced MHC class I downregulation as may be the case for classical MHC class I targets.

Our study has limitations. While Mtb44 was previously shown to bind HLA-E*01:03 with an affinity comparable to the class I signal peptides, our approach failed to identify specific and efficient scFvs for HIV-Gag.275 and SIV-Gag.276, despite previous studies indicating in vivo immunogenicity of these peptide:HLA-E complexes (Yang et al., 2021). Wallace and colleagues (Wallace et al., 2024) on the other hand suggest that HIV-1-derived peptides, including HIV-Gag.275, exhibit insufficient affinity to HLA-E for stable peptide presentation. It is unclear whether phage panning for these targets was unsuccessful due to insufficient library diversity, lack of peptide binding or stochastic factors during the selection process, and further investigation into low-affinity HLA-E binding peptides is warranted to determine the suitability of this approach for other targets. Similarly, we found that RLP-13 efficiently mediated killing of cells expressing the HIV-1 NL4-3 *Δenv*-eGFP construct encoding for Mtb44, but the question remains whether HIV-1-derived target peptides with lower affinity to HLA-E would also be presented on the cell surface at sufficient levels, or whether competition with the high-affinity signal peptides would prevent this. Efforts to identify additional HLA-E-binding peptides in the setting of cancer and infections are ongoing and will undoubtedly further our understanding of the limitations of HLA-E presentation and targeting.

## Methods

### Cell Line Generation

The following primers were used to linearize the pCMMP-MCS-IRES-puro vector (Addgene) (Primer 1 GAGGGCCCGGAAACCTGG, Primer 2 AGCTCTGCTTATATAGACCTCCCAC). A gene block including the HLA-E*01:03 sequence followed by T2A and BFP genes were inserted by Gibson assembly into the linearized vector. Successful cloning was evaluated by colony PCR and Sanger sequencing. The construct was subsequently linearized and nucleofected into wild-type K562 cells using the Lonza 2D nucleofector. Cells were plated in BCM and assessed for BFP expression at three days post nucleofection. Nucleofected cells were then sorted for high BFP expression and these cells were transferred to 1.25 µg/ml puromycin for selection. BFP expressing cells that grew in these cultures were subsequently sorted into single-cells on a 96-well plate with puromycin-containing BCM. Single-cell clones were assessed by flow to evaluate for stable BFP expression. Clone 3.3 was selected as stably containing an elevated level of BFP expression and was further expanded and characterized.

### Phage-Display Library Construction

The scFv-bearing library used for phage panning was previously described (Douglass et al., 2021; Hsiue et al., 2021) and was regrown within a week of selection.

### Phage Panning and Characterization and scDb preparation

Phage panning, characterization, and generation of scDbs were performed as previously described (Sengupta et al., 2022) with a few modifications to allow for HLA-E selection. Briefly, panning consisted of four rounds of negative and positive selection. Negative selection was performed against naked streptavidin beads (FisherScientific, Cat. # 11206D), free streptavidin, and irrelevant, and biotinylated HLA-E-pMHC pre-conjugated to streptavidin beads. Negative selection was also performed against HLA-E-bearing cells (K562-3.3 described above) either un-pulsed or pulsed with various peptides. Negative selection was also performed against HLA-E overexpressing cells bearing the following epitopes HIV.Gag275 (RMYSPTSIL), Mtb44 (RLPAKAPLL), SP-2A (VMAPRTLVL) (kind gift from the group of Prof. Andrew McMichael).

Positive selection was performed for each round of panning with decreasing amounts of the relevant biotinylated monomer conjugated to streptavidin beads. After each round of panning, phage were eluted with glycine (pH 2), neutralized with Tris-HCL (pH 9), amplified in SS320 cells (Lucigen), and concentrated in PEG-NaCl. Phage eluted from the 3rd and 4th round of panning were diluted such that infection of bacteria produced single colonies. Individual bacterial colonies were inoculated in deep 96-well plates to produce monoclonal phage. Monoclonal phage supernatants were tested for their ability to bind to the target versus irrelevant pMHC using ELISA and flow cytometry. Phage supernatants were added to streptavidin ELISA plates (R&D systems) pre-coated with biotinylated HLA-E-monomers (1 µg/mL) of the relevant or irrelevant pMHC, and phage binding was assessed using a rabbit anti-M13 antibody (Pierce) and a secondary conjugated to HRP.

To assess phage binding to pMHC on biotinylated plates, T2 cells were pulsed with 50 µg/ml peptide and 10 µg/ml B2M for 4 hours to overnight at 37°C in RPMI 1640 + Glutamax with 1% Pen/Strep. Pulsed cells were washed once with PBS, incubated with monoclonal phage, and then stained with rabbit anti-M13 (Pierce) followed by a PE-conjugated donkey anti-rabbit secondary (Biolegend). Phage-stained cells were acquired on the Intellicyt flow cytometer (Satorius) and analyzed with FlowJo (v. 10.10.0).

### scDb Expression and Purification

Monoclonal phages that stained specifically for a particular peptide-pulsed target were sequenced to identify the expressed Fab. Gene blocks expressing this Fab fragment, the UCHT1 Fab fragment against CD3, and C-terminal His tag were cloned via Gibson assembly into the pcDNA3.4 vector backbone. scDb constructs were sequenced, amplified, and expressed via transfection into HEK 293T cells. scDbs were purified using nickel columns (Capturem His-tag miniprep columns, TakaraBio) using 400 mM imidazole and were desalted to remove imidazole using Zeba Spin columns (7K MWCO, FisherScientific). scDbs were run on SDS-PAGE and quantified using iBright densitometric analysis in comparison to a BSA standard. Large-scale preparations of scDbs were produced by ThermoFisher.

### ScDb binding ELISA

Streptavidin-coated plates (R&D Systems) were incubated overnight with biotinylated peptide-MHC monomers at 1 µg/mL, followed by three washes with PBS-Tween. ScDb was added at a concentration of 5 µg/mL in 50 µL BAE buffer (PBS + 0.5% BSA + 0.1% sodium azide) for 1 hour at room temperature. After three washes, HRP anti-6x His tag mIgG secondary (clone J099B12, BioLegend, 0.5 ug/ml in BAE) was added for another 1 hour incubation at room temperature. Plates were washed an additional 3 times and 50 uL per well of TMB substrate (BioLegend) was added. The reaction was quenched with 1N sulfuric acid and absorbance at 450 nM was read using the VarioSkan plate reader (FisherScientific).

### Viral constructs and virus generation

For the generation of an Mtb44-expressing viral reporter construct based on an existing NL4.3 ΔEnv-eGFP backbone, the DNA sequence encoding for the Mtb44 peptide (5’-CGCCTTCCAGCAAAAGCTCCTCTGCTG-3’) was inserted at the 3’-end of the eGFP open reading frame before the STOP codon. A T2A site (5’-GAAGGCAGAGGCTCTCTGCTTACATGTGGCGACGTGGAAGAGAACCCCGGACCT-3’) was included between the eGFP and Mtb44 coding sequences to prevent Mtb44 from impacting eGFP function. Cloning was performed by GenScript. Correct insertion was confirmed by Sanger sequencing.

Virus was generated by co-transfecting HEK 293T cells with plasmids encoding for backbone (NL4.3 ΔEnv-eGFP or NL4.3 deltaEnv-eGFP-Mtb44) and HIV Env (Strain 89.6) using the Lipofectamine 3000 Transfection Reagent kit (ThermoScientific, Cat. L3000001) according to manufacturer instructions. The NL4.3 ΔEnv-eGFP backbone expression plasmid was obtained through the NIH HIV Reagent Program, Division of AIDS, NIAID, NIH: Human Immunodeficiency Virus 1 (HIV-1) NL4-3 ΔEnv EGFP Reporter Vector, ARP-11100, contributed by Dr. Haili Zhang, Dr. Yan Zhou and Dr. Robert Siliciano. The HIV env expression vector was obtained through the NIH HIV Reagent Program, Division of AIDS, NIAID, NIH: Human Immunodeficiency Virus-1 89.6 Env Expression Vector (pcDNA 89.6 env), ARP-12485, contributed by Dr. Kathleen Collins and Dr. Ronald Collman. Virus-containing supernatant was harvested 48 hours and 72 hours post-transfection, and virus was isolated using a 0.22 µm filter and ultracentrifugation with 20% sucrose. Isolated virus was then titered using a p24 ELISA kit (PerkinElmer) following the manufacturer protocol.

### Cell culture

All cells were kept in incubators at 37°C with 5% CO2. Cells derived from K562 cell lines were cultured in IMDM (Gibco, Cat. 12440) with 8% heat-inactivated fetal bovine serum (GeminiBio) and 1% Penicillin/Streptomycin (ThermoFisher). Peripheral mononuclear cells (PBMCs) from HIV-seronegative donors were obtained from StemCell and viably frozen in fetal bovine serum with 10% DMSO (SigmaAldrich) at a cell density of 1*10^8^ cells/mL to be stored in liquid nitrogen until use. PBMCs were thawed in base media (RPMI1640 + GlutaMAX (Gibco, Cat. # 61870) with 10% fetal bovine serum and 1% Penicillin/Streptomycin) and rested for 3-24 hours before further use. CD8^+^ T cells were prepared by stimulating PBMCs with 15 ng/mL of anti-CD3 mAb (BioLegend, Cat. # 317302) in CD8 media (base media + 100U/mL recombinant human IL-2 (R&D Systems, Cat. # 202-IL-050/CF) + 5 ng/mL recombinant human IL-7 (BioLegend, Cat. # 581906)) at a starting cell density of 2*10^6^ cell/mL for 72 hours. PBMCs were then washed and resuspended in CD8 media for 72 hours before isolation of CD8^+^ T cells using a CD8 T cell Enrichment kit (StemCell, Cat. # 19053) following the manufacturer protocol.

CD8^+^ T cells were rested for an additional 4 days in CD8 media before co-culture setup. CD4^+^ T cells were isolated from unactivated PBMCs using the CD4 T Cell Enrichment kit (StemCell, Cat. # 19052) following the manufacturer protocol. CD4^+^ T cells were then activated using ⍺CD3/⍺CD28 T cell activator Dynabeads (FisherScientific, Cat. # 11131D) at a bead-to-cell ratio of 1:1 and a cell density of 1*10^6^ cells/mL in CD4 media (base media + 30U/mL rhIL-2) for 72 hours before infection. Cells were infected by centrifugation of 1*10^5^ cells per well in 96-well flat-bottom plates with the indicated HIV-1 reporter constructs (300 ng p24/1*10^5^ cells) for 2 hours at 1,200x g and 30°C and cultured in T Cell Growth Media (RPMI-1640 with GlutaMAX, 20 U/mL penicillin, 20 µg/mL streptomycin, 10% heat-inactivated fetal bovine serum, 1% T-cell growth factor (Siliciano and Siliciano, 2005), 100 U/mL recombinant human IL-2) for 72 hours before the co-culture. HEK293T cells were cultured in DMEM + GlutaMAX (Gibco, Cat. # 10569) with 10% FBS and 1% Penicillin/Streptomycin. Cells derived from the K562 cell line were cultured in IMDM (Gibco, Cat. # 12440053) with 8% FBS and 1% Penicillin/Streptomycin. For peptide pulsing, cells were cultured for 4 hours in low-serum media (K562: IMDM + 1% Penicillin/Streptomycin, PBMC: RPMI-1640 with Glutamax + 2% FBS + 1% Penicillin/Streptomycin) at 37°C.

### Co-culture setup

PBMCs from HIV-negative donors were stimulated with soluble anti-CD3 monoclonal antibodies (BioLegend, clone OKT3, 15ng/mL, Cat. # 317302) for 72 hours in CD8 media (RPMI1640 (FisherScientific)+ 10% Fetal Bovine Serum (GeminiBio) + 1% Penicillin/Streptomycin (ThermoFisher) + 100 U/mL IL-2 (R&D Systems) + 5 ng/mL hrIL-7 (BioLegend)). Anti-CD3 mAbs were then washed out, and CD8^+^ T cells were isolated using the CD8 T cell Enrichment kit (StemCell, Cat. #19053). CD8^+^ T cells were cultured without stimulation for 7 days before the killing assay.

For our dual-color co-culture (Figure 4), pulsed cells were incubated for 15 minutes in PBS with CellTrace Violet or CFSE (FisherScientific, Cat. # C34557) or CFSE (FisherScientific, Cat. # C34554) at a 1:20,000 dilution. Cells were then washed and resuspended in culture media and added to the co-culture plate with primary CD8^+^ T cells and with RLP-13 scDb or irrelevant control H2-mu scDb at an effector:cognate:non-cognate target ratio of 4:1:3. After the co-culture, supernatant was saved to be used in cytokine quantification assays using the LegendPlex CD8/NK Cell Panel kit (BioLegend, Cat. # 741187) following the manufacturer protocol.

For peptide titration co-cultures (Figure 5), PBMCs from HIV-negative donors were pulsed with peptide at the indicated concentrations for 4 hours in low-serum media and co-cultured with autologous CD8^+^ T cells and with or without RLP-13 scDb at an effector-to-target ratio of 3:1. Supernatant was used for cytokine quantification as described above.

For primary cell co-cultures with infected cells(Figure 6), PBMCs from HIV-negative donor PBMCs were activated for 72 hours in T Cell Growth Media with PHA (0.5 µg/mL). After 72 hours, CD4^+^ T cells were isolated through negative selection (StemCell, CD4 T Cell Enrichment Kit) and infected with the indicated viral construct (300 ng p24 per 1*10^5^ cells) by centrifugation of 1*10^5^ cells per well in a 96-well flat-bottom plate in T Cell Growth Media with virus in a total volume of 40 µL. After centrifugation, cells were moved to an incubator at 37°C and 5% CO2 to rest for 3 hours. Subsequently, all individual wells from the same condition were pooled, and cells were washed and resuspended at a density of 0.5 M/mL in T Cell Growth Media. After 72 hours, the cells were co-cultured for 18 hours at 37°C with primary CD8^+^ T cells at an effector-to-target ratio of 1:1 and with or without RLP-13 scDb. T cell activation and target cell viability were assessed through surface staining with anti-CD8 (BV605, clone SK1), anti-CD3 (BV421, clone OKT3), anti-CD69 (APC, clone FN50) (all BioLegend) and eBioscience Fixable Viability Dye eFluor780 (FisherScientific) and analysis by flow cytometry. HIV-1-infected cells were identified through the eGFP reporter among CD3^+^CD8^-^ cells (Figure S5). Target cell viability for each condition was normalized to the target cell viability in the absence of scDb.

### Flow cytometry

Cells were centrifuged at 400x g for 5 minutes. Supernatant was discarded and cells were washed in 200uL Phosphate-buffered Saline (PBS, FisherScientific, Cat. # 10010) and centrifuged again at 400x g for 5 minutes. Cells were then stained in 50 µL of the respective staining mix in PBS. All antibodies are obtained from BioLegend and at a 1:70 dilution unless noted otherwise: APC-conjugated anti-human CD69 (clone FN50, Cat. No# 310910), Brilliant Violet 421-conjugated anti-human CD3 (clone OKT3, Cat. # 317344), Brilliant Violet 605-conjugated anti-human CD8 (clone SK1, Cat. # 344742), PE-conjugated anti-human HLA-A2 (Cat. No# 343306), PE-conjugated anti-human HLA-E (Cat. # 342604), Brillian Violet 421-conjugated anti-human CXCR4 (Cat. # 306518). We also used eBioscience 780nM Fixable Viability Dye (1:1000, FisherScientific, Cat. # 65-0865-14). For dual-color co-cultures, target cells were stained with CellTrace Violet or CFSE (dilution 1:20,000, FisherScientific, Cat. # C34557 and Cat # C34554) for 15 minutes in PBS at 37°C prior to plating. Co-cultures from Figure 4 and 5 were run on the Satorius Intellicyt iQue Screener Plus flow cytometer, and co-cultures from Figure 6 were run on the Cytek Northern Lights (3L V-B-R configuration). All flow cytometry data was analyzed using FlowJo v.10.10.0.

### Materials

Biotinylated HLA-E*01:03/Mtb44, HLA-E*01:03/HIV.Gag275, HLA-E*01:03/SIV.Gag276, HLA-E*01:03/SP-2A and other monomers were obtained from the Baylor College of Medicine MHC Tetramer Core Facility. Peptides were obtained from Elim Biopharm at a purity of >95%.

### Surface plasmon resonance (SPR) affinity measurements

Binding kinetics of RLP-13 to Mtb44/HLA-E*01:03, HIV-Gag.275/HLA-E*01:03, and SP-2A/HLA-E*01:03 were measured using a Biacore T200 (Cytiva) with SA chip at 25°C. Biotinylated Mtb44/HLA-E*01:03 (1 mg/ml stock), HIV-Gag.275/HLA-E*01:03 (1 mg/ml stock), and SP-2A/HLA-E*01:03 (1 mg/ml stock) were used as ligands to capture on the SA sensor surface. RLP-13 (17.9 μM stock) was injected as the analyte. The ligands were diluted to 20 ng/ml in HBS-P (10 mM Hepes pH 7.4, 150 mM NaCl, 0.05% v/v surfactant P20) and each captured onto a different Flow Cell to a level of ∼30 Response Units (RU). Flow Cell 1 with no captured ligand was used as a reference. HBS-P was used as the immobilization capture running buffer. Based on the ligand captured response values, the theoretical Rmax values for each analyte were about 35.5 RU, assuming a 1:1 interaction mechanism. Overnight kinetics were performed for the analytes binding to the captured ligands in the presence of HBS-P supplemented with 5% Glycerol. The flow rate of all analyte solutions was maintained at 50 μL/min. Injected analyte concentrations were from 100 nM to 3.125 nM (two-fold dilutions) for multicycle kinetics. All analytes were injected in duplicate. The association phase was monitored for 120 s, followed by a dissociation time of 600 s. All analyte injections were performed in duplicate. Regeneration of the sensor surface was achieved with a single 20-s injection of 2 M NaCl. Binding kinetics were analyzed by fitting the data to a two-state reaction model using Biacore evaluation software.

## Data availability

All relevant data are available within the paper and supplementary materials, and upon request from the lead contact, R.F. Siliciano. Further information and requests for reagents generated or used in this study are also available upon request from the lead contact.

## Acknowledgements

This work was supported in part by The Bloomberg-Kimmel Institute for Cancer Immunotherapy. The authors acknowledge the use of the Biacore Molecular Interaction Shared Resource (BMISR) at Georgetown University, which is supported by the National Institutes of Health (P30CA51008). Biotinylated peptide-bearing HLA-E*01:03 monomers were obtained from the Baylor College of Medicine MHC Tetramer Core Facility.

## Author contributions

S. Sengupta and R.F. Siliciano conceived the study. S. Sengupta, N. Bachmann, J. Douglass, N. L. Board, F. Wu and M. Moskovljevic developed the methods. J. Douglass and S. DiNapoli generated phage library. S. Sengupta and N. Bachmann performed phage panning, selection, and *in vitro* experiments with assistance from J. Zhang, N. L. Board, F. Wu, and M. Moskovljevic. Y. Tabdili, M. Reed, and J. Yu assisted with the generation of HLA-E expressing cell lines. P. A. Azurmendi designed and performed SPR experiments and analysis. B. Kim assisted with experiments and data acquisition. H. Zhang performed cell sorting. S. Sengupta, N. Bachmann, J. Zhang, J. Douglass, N. L. Board, F. Wu, M. Moskovljevic, S. DiNapoli, Y. Tabdili, M. Reed, J. Yu, P. A. Azurmendi, B. Kim, S. B. Gabelli, J.D. Siliciano, and R. F. Siliciano assisted with analysis and interpretation of the data. S. Sengupta and N. Bachmann wrote the original draft. R. F. Siliciano reviewed and edited the final manuscript. S. B. Gabelli, J. D. Siliciano and R. F. Siliciano contributed reagents/analytic tools. J. D. Siliciano and R. F. Siliciano supervised the study.

---

**Supplementary Figure 1:**
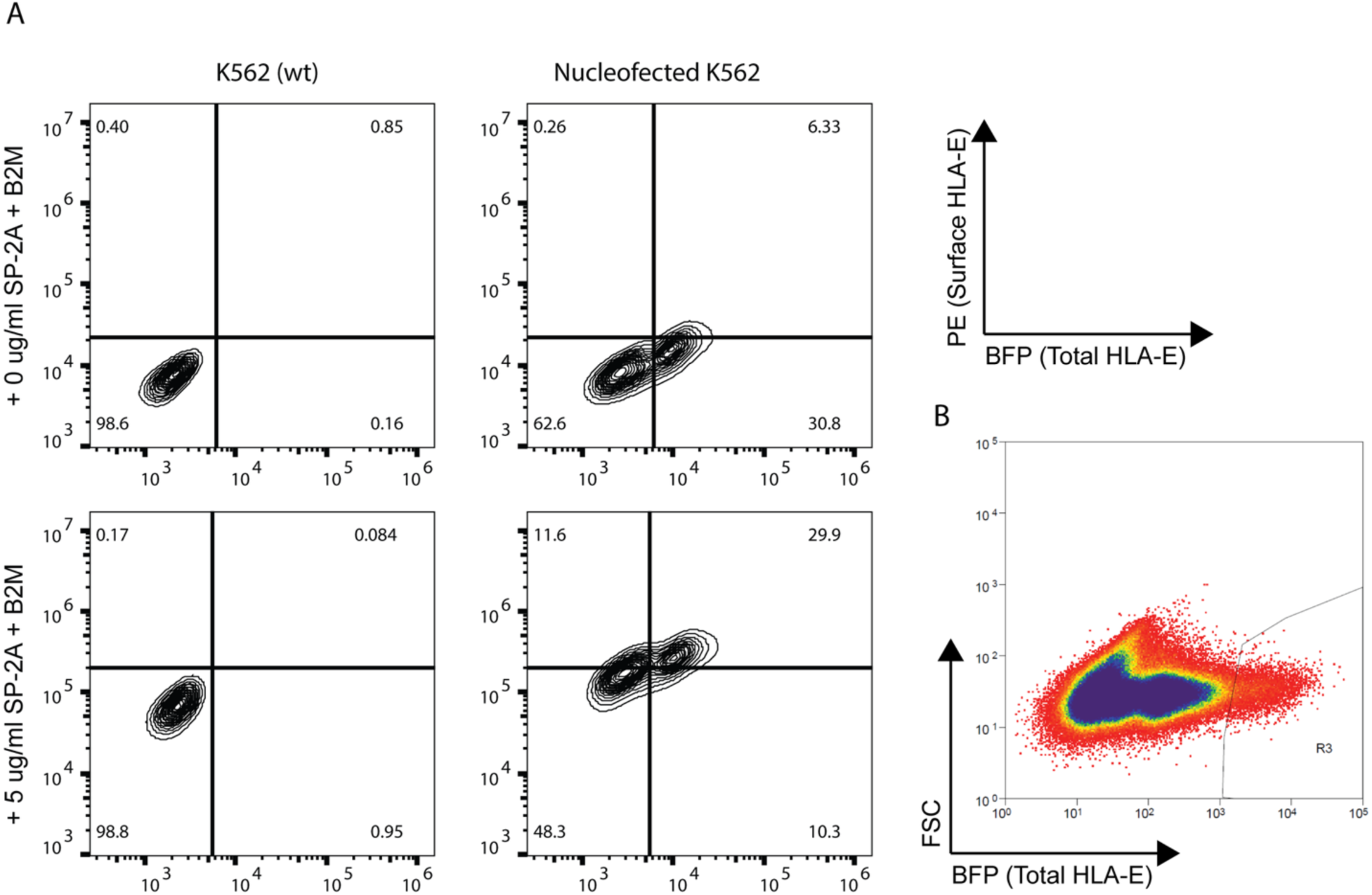
Generation of an HLA-overexpressing K562-derived cell line. (A) Comparison of HLA-E expression between K562 wild-type cells and K562 cells nucleofected with an BFG-tagged HLA-E expression vector. Both cell lines were pulsed with either beta-2-microglobulin (B2M) only or in addition to 5ug/mL SP-2A peptide. Surface levels of HLA-E were assessed through staining with PE-conjugated anti-HLA-E monoclonal antibody (clone 3D12) and measured by flow cytometry. (B) Cell sorting for highly HLA-E-overexpressing cells. HLA-E expression was assessed through BFP-signal, and viable cells that fell into the indicated gate (R3) were retained for further selection.

**Supplementary Figure 2:**
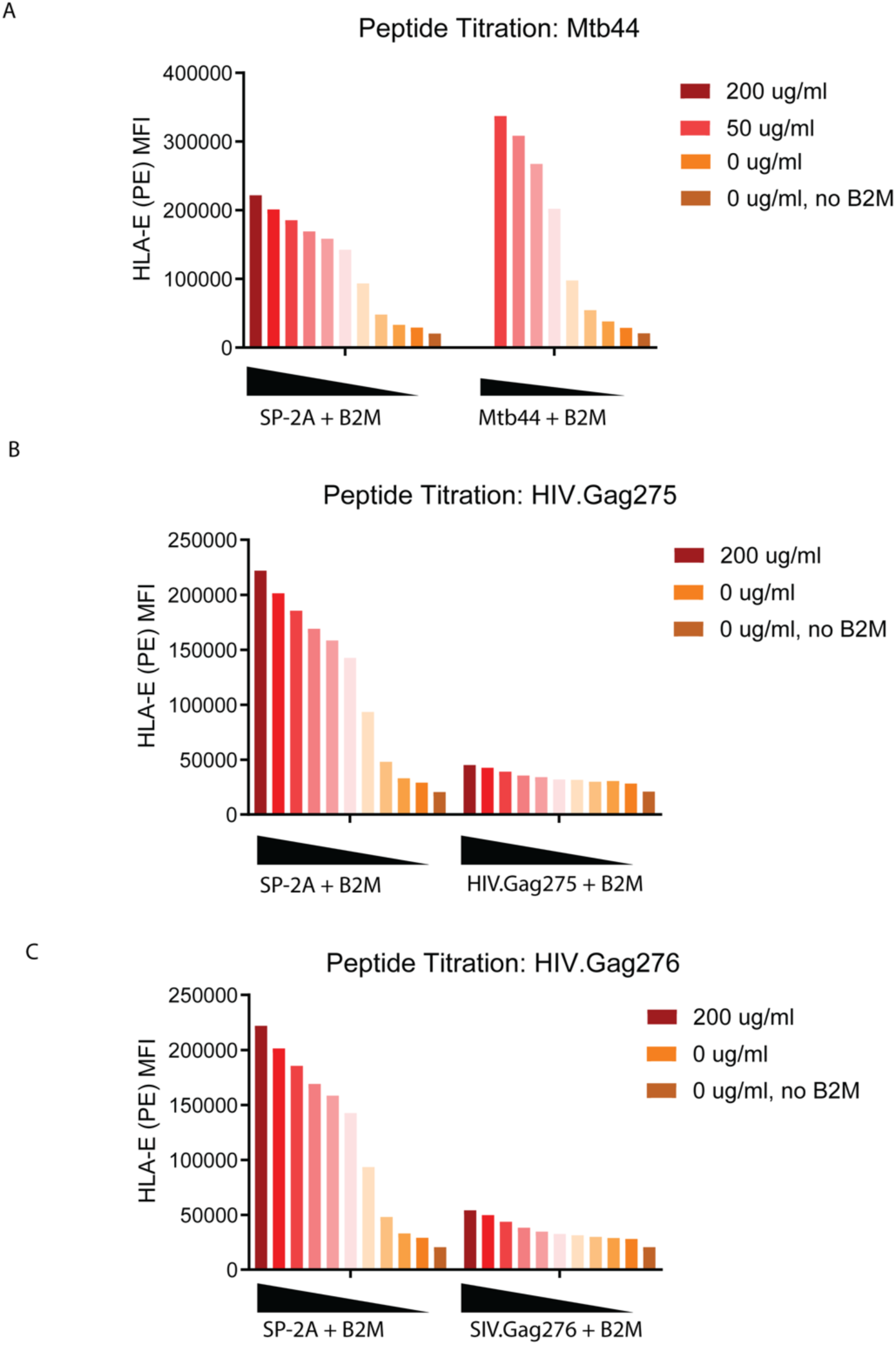
Stabilization of HLA-E on the surface of K562-3.3 cells pulsed with the canonical ligand SP-2A compared to (A) Mtb44, (B) HIV.Gag275 and (C) SIV.Gag276. Cells were cultured in serum-free media with B2M and varying concentrations of indicated peptides in 2-fold serial dilution for 4 hours before staining with PE-conjugated anti-HLA-E monoclonal antibodies and measured by flow cytometry.

**Supplementary Figure 3:**
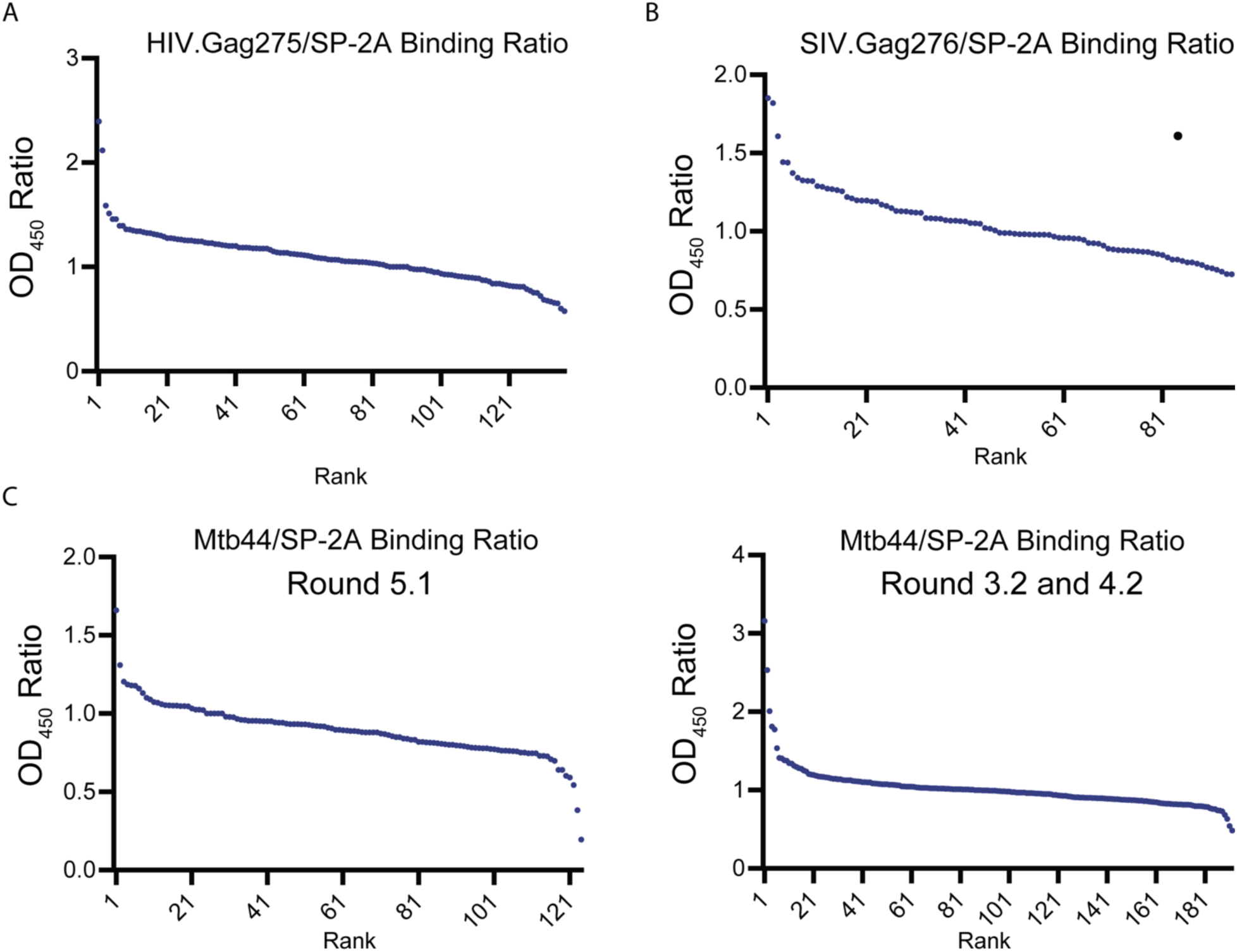
Signal ratios (OD_450_) from pMHC-binding ELISA for isolated phage isolates after 5 rounds of panning for the indicated pMHC target. HLA-E monomers loaded with (A) HIV.Gag275, (B) SIV.Gag276 or (C) Mtb44 were immobilized on streptavidin-coated wells and incubated with monoclonal phage isolates, followed by incubation with HRP-conjugated anti-M13 phage antibody. TMB substrate was added and OD450 was read out and normalized to a matched well with immobilized SP-2A/HLA-E monomer on the same plate. For (C), additional panning was performed with a more stringent protocol starting with the eluate from round 2 of the initial panning, resulting in rounds 3.2 and 4.2. OD_450_ ratios for clones from these rounds are shown on the right. Each dot represents one tested phage clone.

**Supplementary Figure 4:**
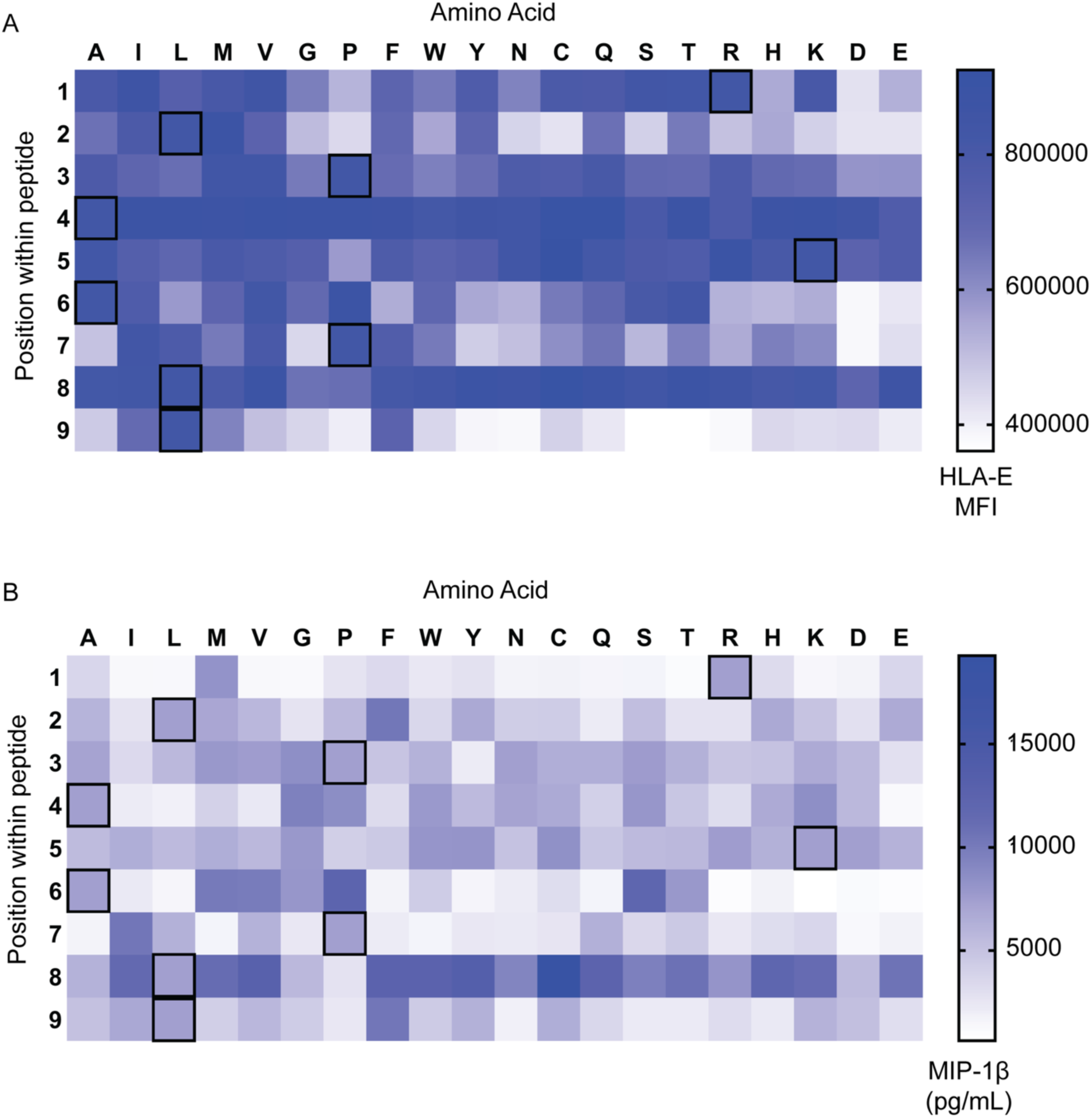
Impact of single residue substitution in Mtb44 on the interactions with HLA-E and RLP-13. K562-3.3 Cells were pulsed with a library of 171 variant peptides where each residue (rows) was substituted with each other possible amino acid (columns) for 4 hours in serum-free media. (A) Cells were stained with PE-conjugated anti-HLA-E monoclonal antibody and surface levels of HLA-E were assessed through flow cytometry. (B) PBMCs were stimulated with anti-human CD3 monoclonal antibody and CD8-positive T cells were isolated and cultured without activating stimuli for 7 days. Pulsed cells from (A) were co-cultured with CD8^+^ T cells and RLP-13. The concentration of chemokine MIP-1β in the co-culture supernatant was quantified through ELISA. The framed cells in the heatmaps correspond do the amino acid sequence in the original Mtb44 peptide.

**Supplementary Figure 5:**
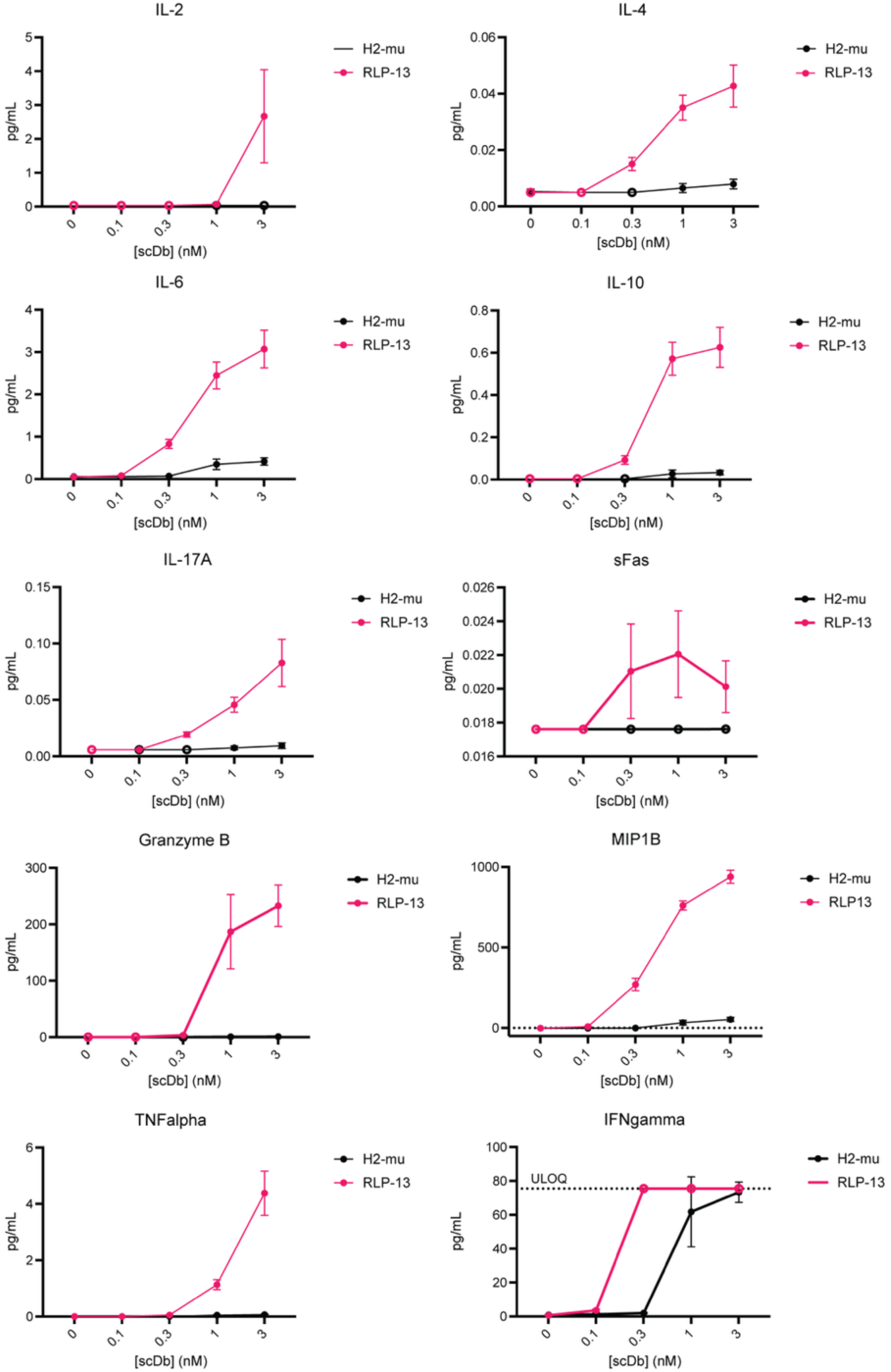
Quantification of additional secreted cytokines, chemokines and effector molecules in co-culture supernatant from Figure 4C. Quantification performed using the BioLegend LEGENDPlex^TM^ Human CD8/NK Panel (13-plex) kit following manufacturer protocol. ULOQ, upper limit of quantification. Black symbols: supernatant from co-culture with irrelevant control scDb; red symbols: co-culture with RLP-13. Empty symbols: values outside of quantifiable range (either below limit of detection or above ULOQ).

**Supplementary Figure 6:**
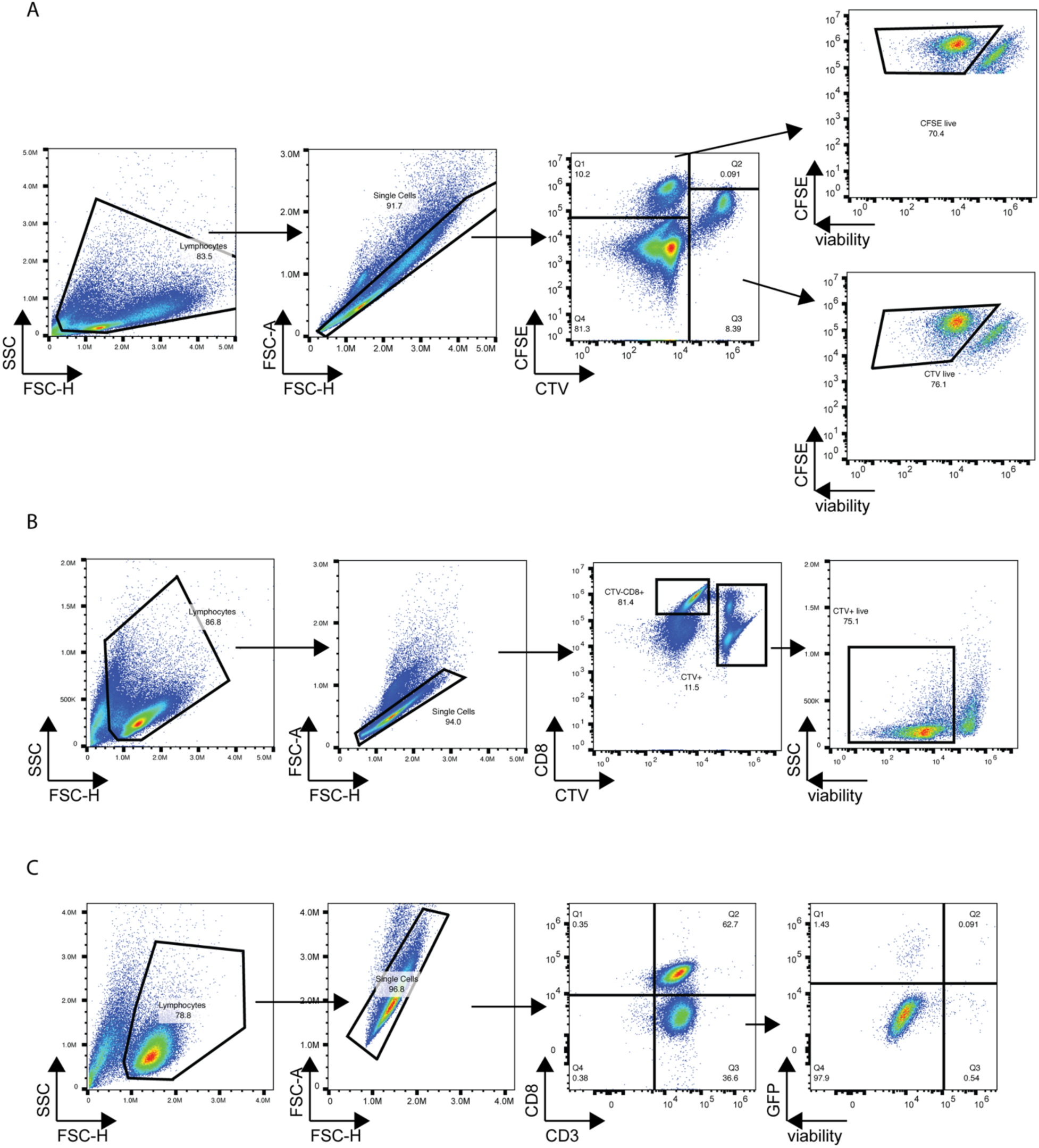
Gating strategies for flow cytometry analysis of co-cultures. (A) Gating strategy for co-cultures in Figure 4C. (B) Gating strategy for co-cultures in Figure 5B. (C) Gating strategy for co-cultures in Figure 6D.

**Supplementary Table 1:**
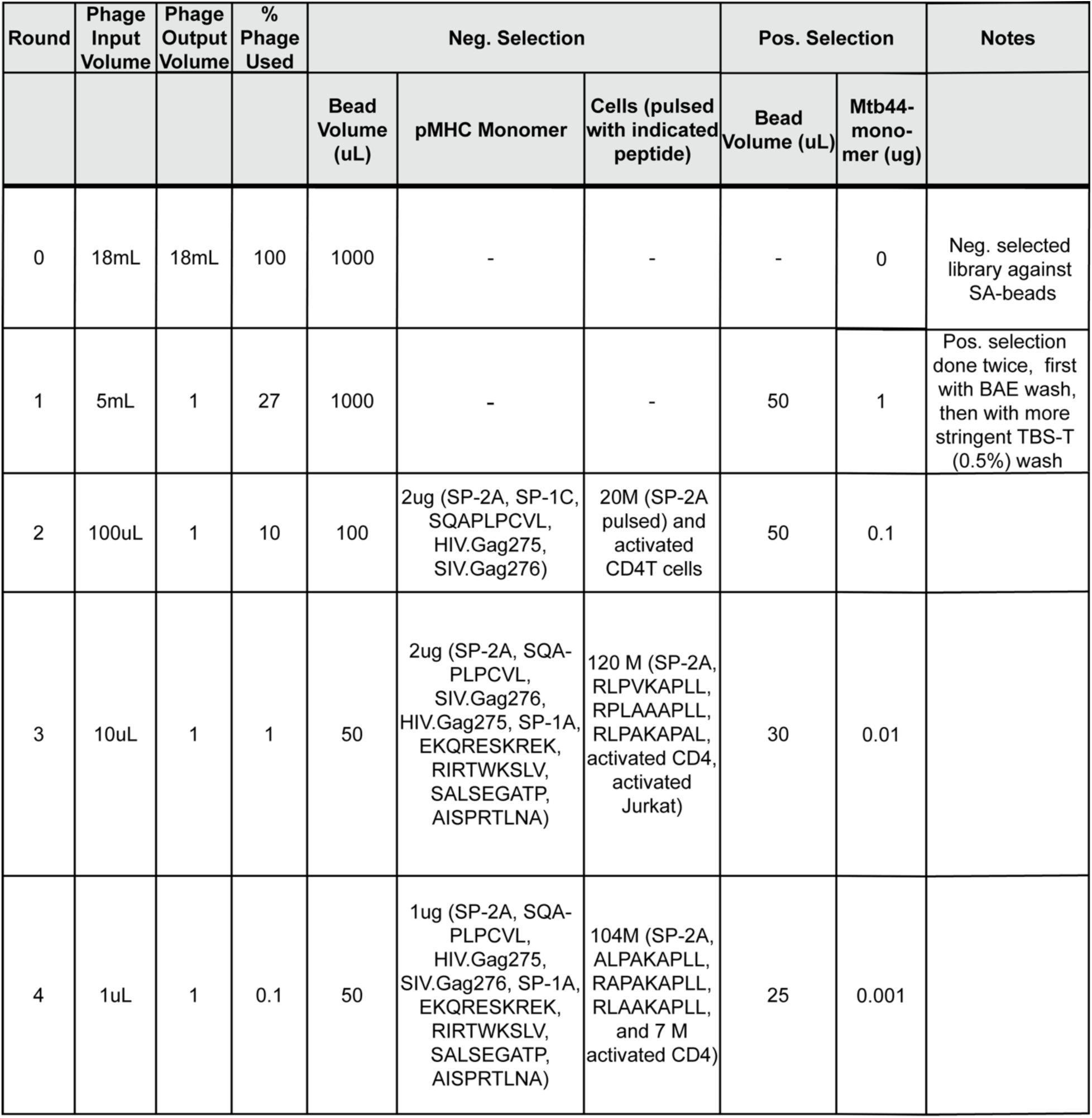
Phage panning conditions for Mtb44/HLA-E*01:03 using the previously published Ludwig3 phage library.

**Supplementary Table 2:**
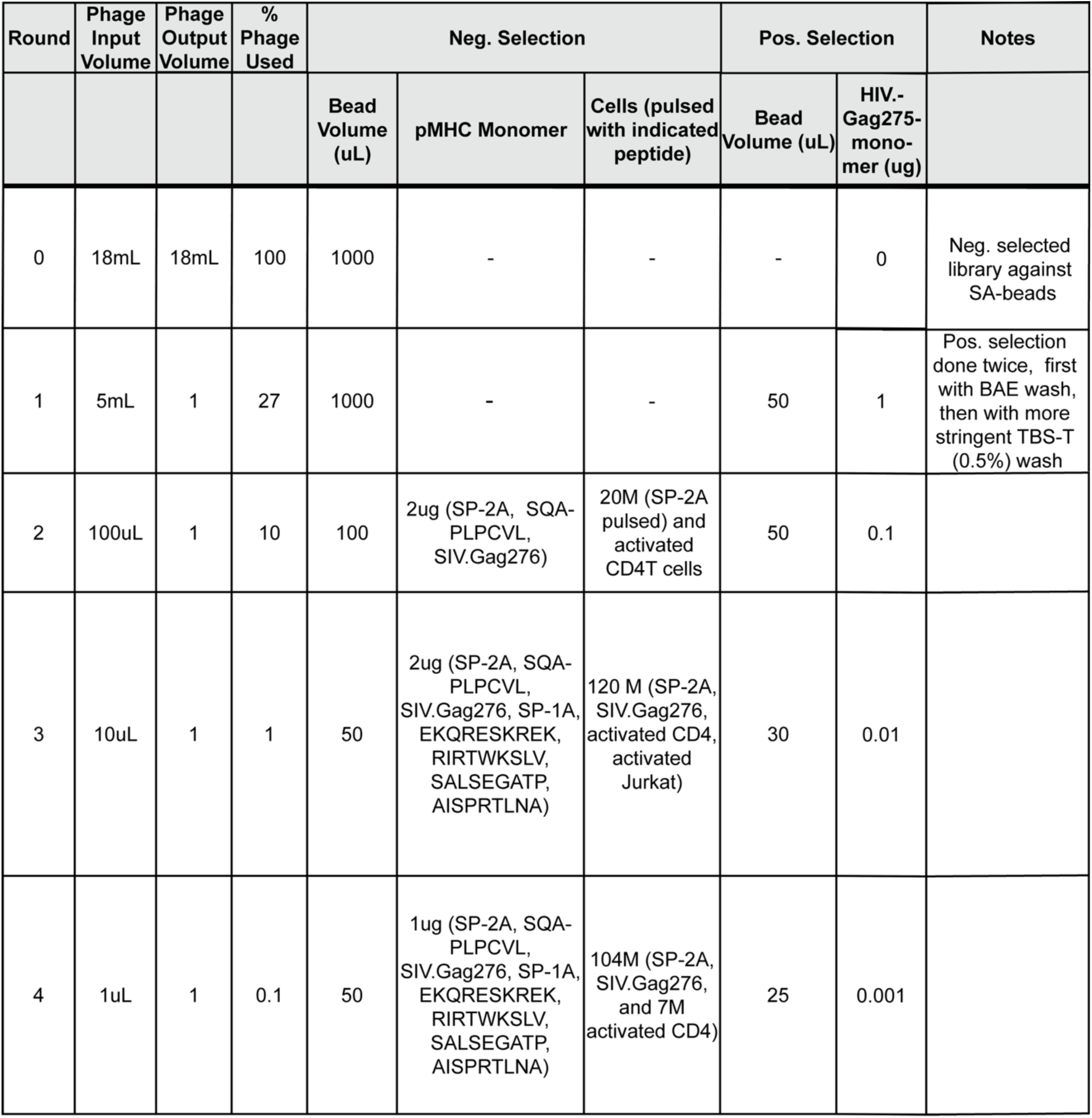
Phage panning conditions for HIV.Gag275/HLA-E*01:03 using the previously published Ludwig3 phage library.

**Supplementary Table 3:**
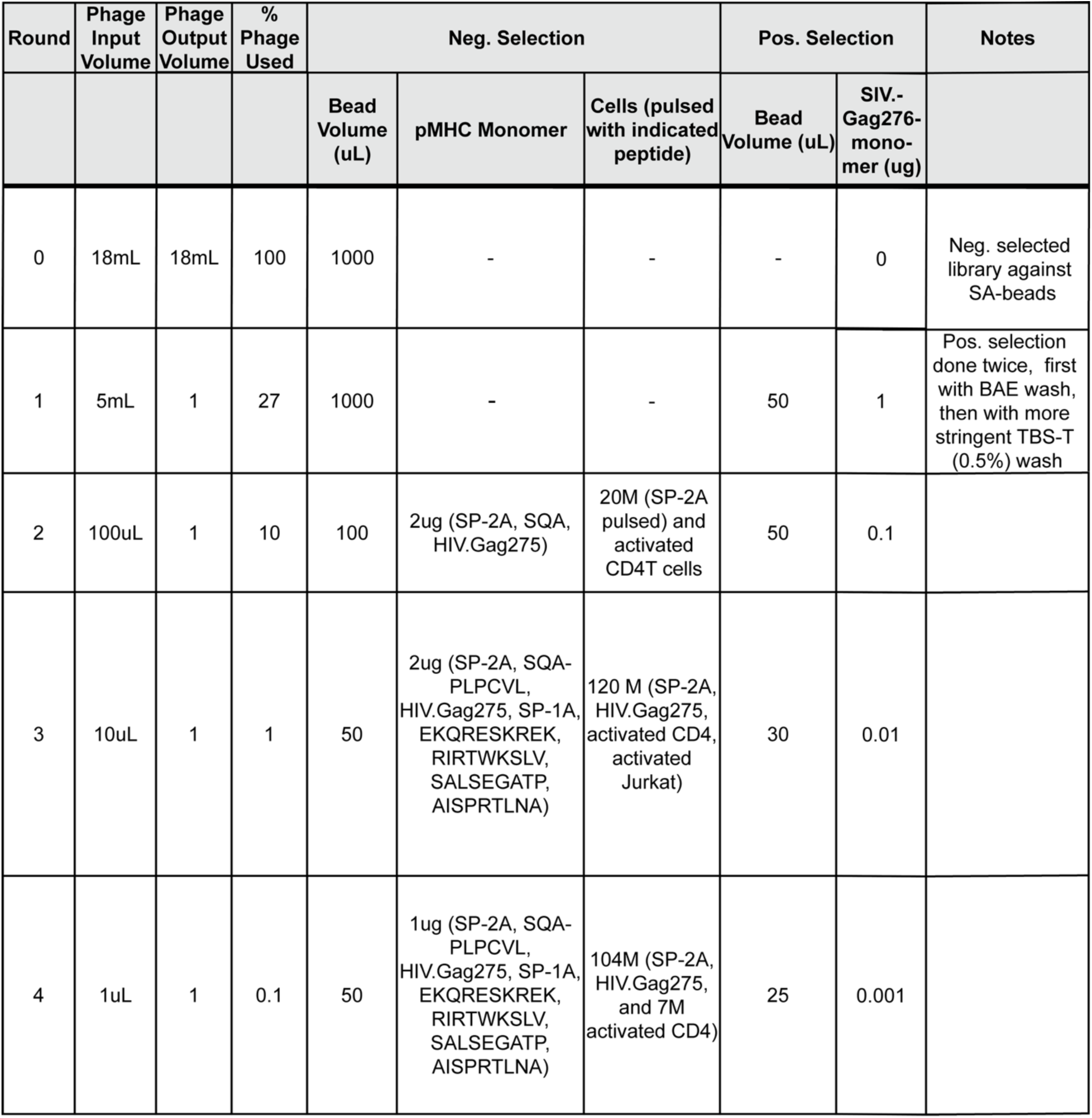
Phage panning conditions for SIV.Gag276/HLA-E*01:03 using the previously published Ludwig3 phage library.

