## Supplementary data for "Potential of HLA-E-targeting diabodies to induce lysis of HIV-1-infected cells by CD8^+^ T cells"

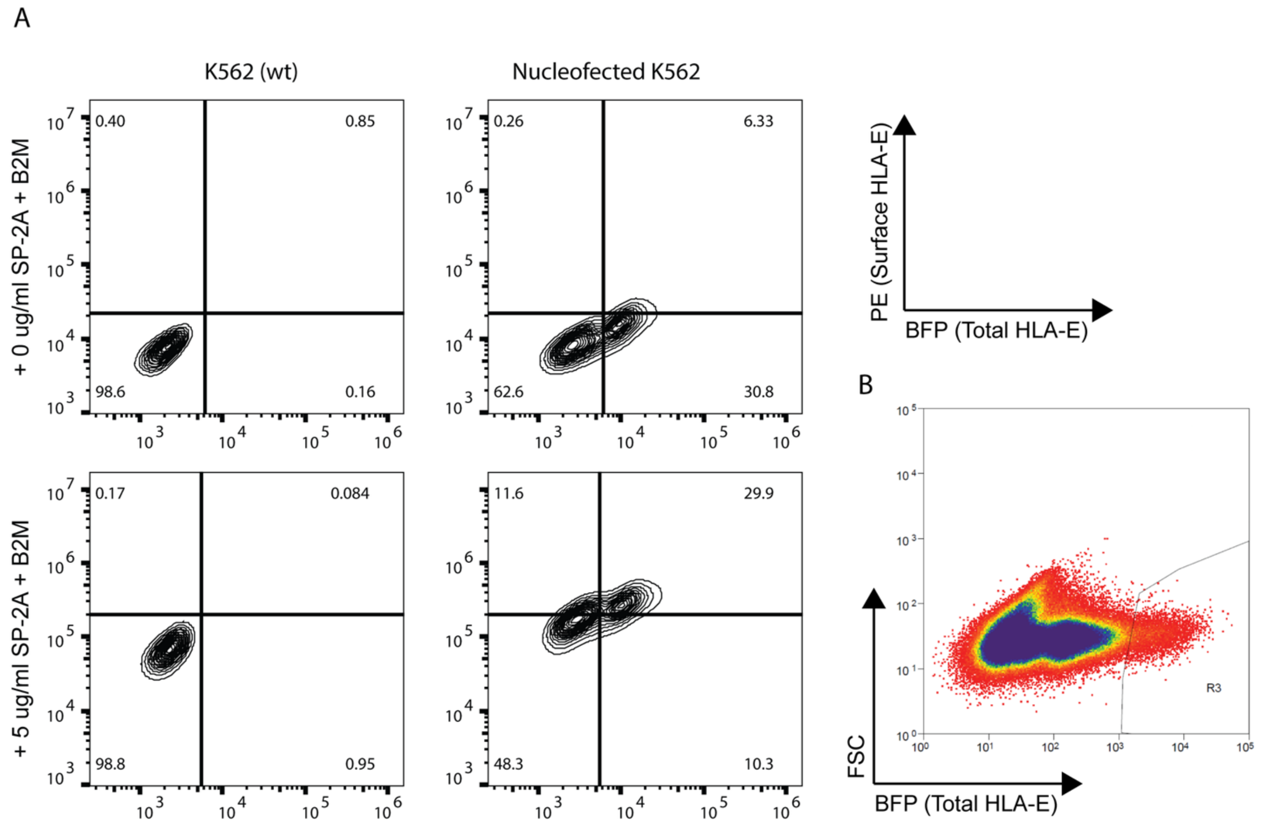


Supplementary Figure 1: Generation of an HLA-overexpressing K562-derived cell line. (A) Comparison of HLA-E expression between K562 wild-type cells and K562 cells nucleofected with an BFG-tagged HLA-E expression vector. Both cell lines were pulsed with either beta-2-microglobulin (B2M) only or in addition to 5ug/mL SP-2A peptide. Surface levels of HLA-E were assessed through staining with PE-conjugated anti-HLA-E monoclonal antibody (clone 3D12) and measured by flow cytometry. (B) Cell sorting for highly HLA-E-overexpressing cells. HLA-E expression was assessed through BFP-signal, and viable cells that fell into the indicated gate (R3) were retained for further selection.

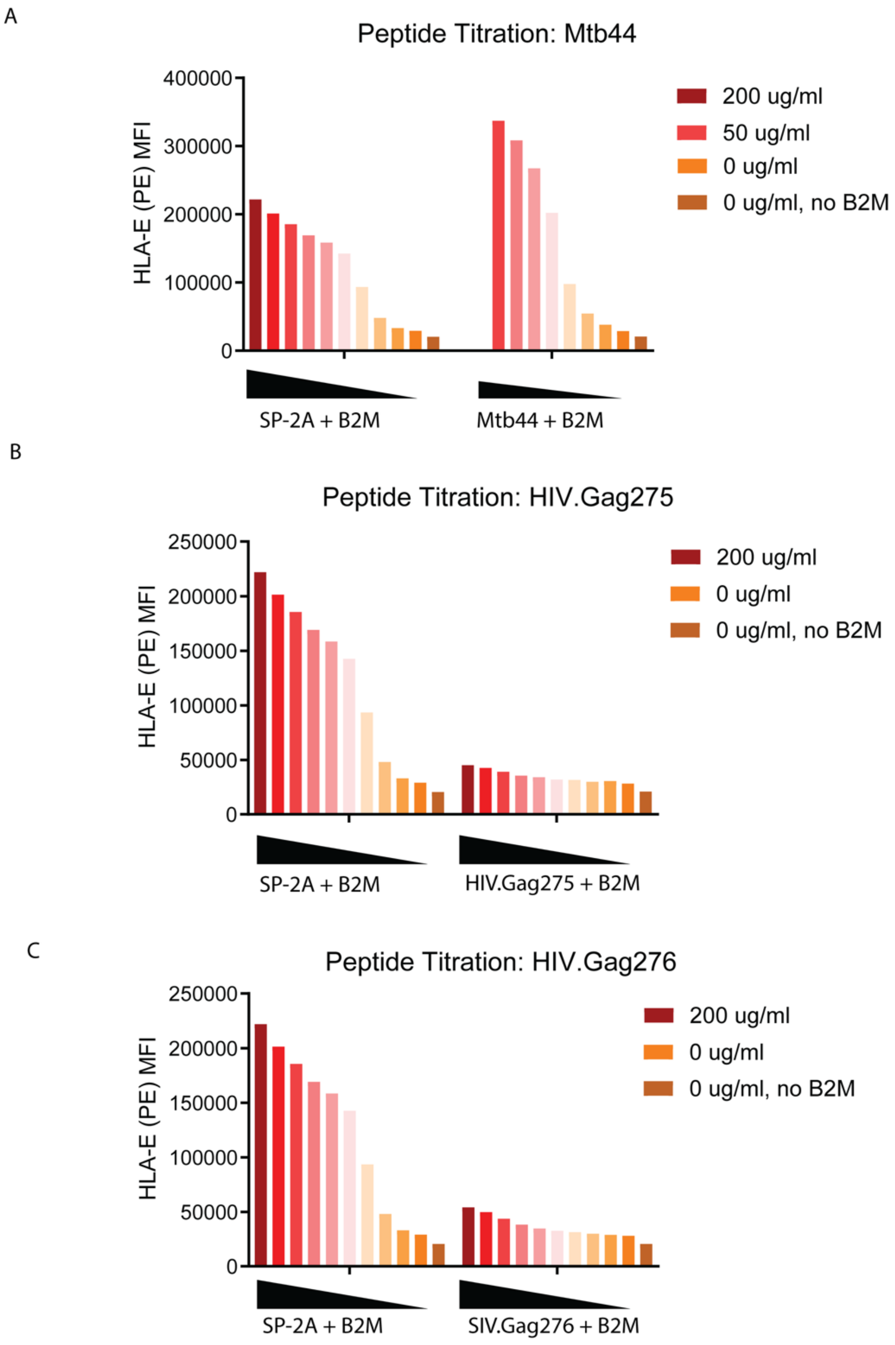


Supplementary Figure 2: Stabilization of HLA-E on the surface of K562-3.3 cells pulsed with the canonical ligand SP-2A compared to (A) Mtb44, (B) HIV.Gag275 and (C) SIV.Gag276. Cells were cultured in serum-free media with B2M and varying concentrations of indicated peptides in 2-fold serial dilution for 4 hours before staining with PE-conjugated anti-HLA-E monoclonal antibodies and measured by flow cytometry.

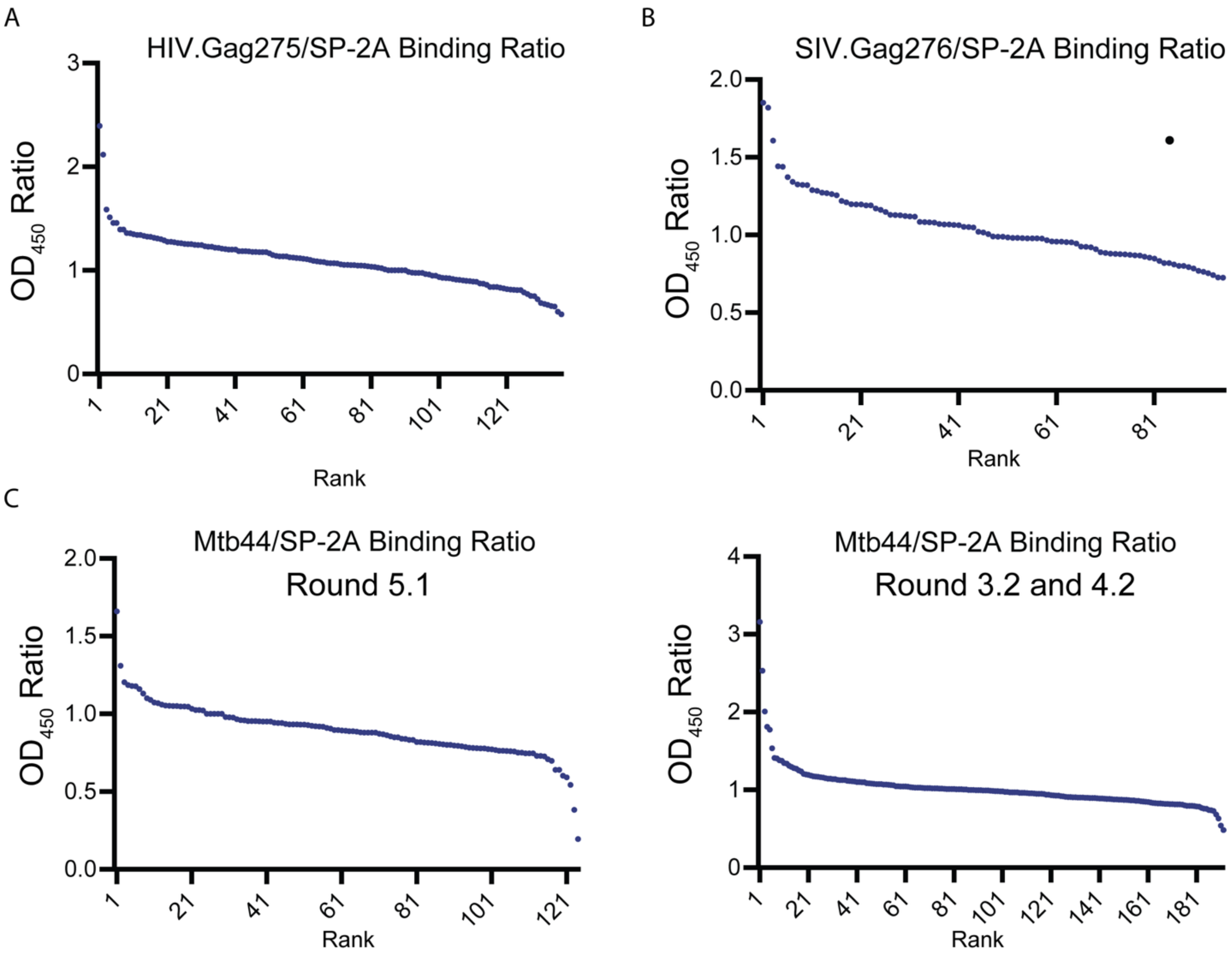


Supplementary Figure 3: Signal ratios ($OD_{450}$) from pMHC-binding ELISA for isolated phage isolates after 5 rounds of panning for the indicated pMHC target. HLA-E monomers loaded with (A) HIV.Gag275, (B) SIV.Gag276 or (C) Mtb44 were immobilized on streptavidin-coated wells and incubated with monoclonal phage isolates, followed by incubation with HRP-conjugated anti-M13 phage antibody. TMB substrate was added and OD450 was read out and normalized to a matched well with immobilized SP-2A/HLA-E monomer on the same plate. For (C), additional panning was performed with a more stringent protocol starting with the eluate from round 2 of the initial panning, resulting in rounds 3.2 and 4.2. $OD_{450}$ ratios for clones from these rounds are shown on the right. Each dot represents one tested phage clone.

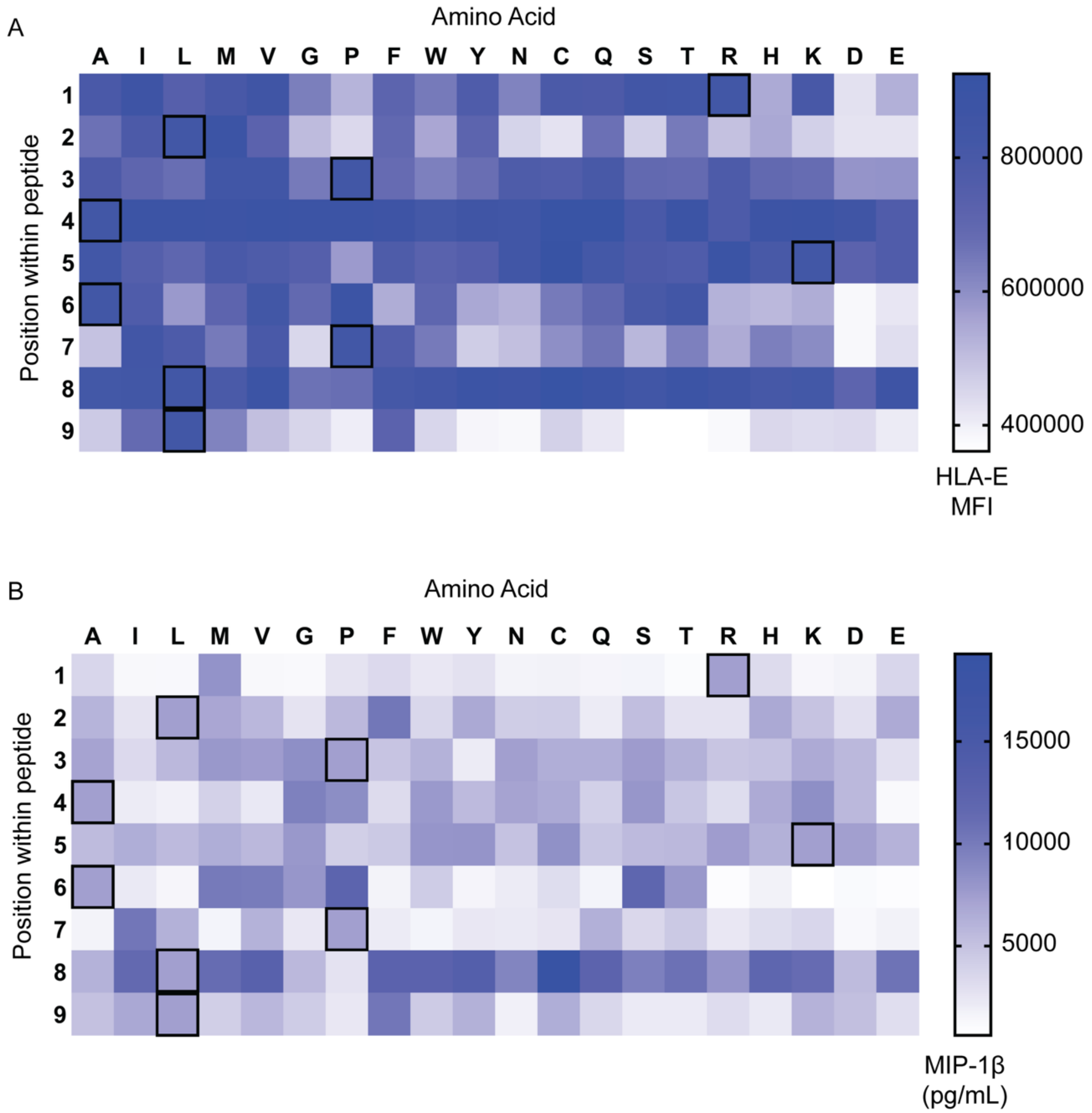


Supplementary Figure 4: Impact of single residue substitution in Mtb44 on the interactions with HLA-E and RLP-13. K562-3.3 Cells were pulsed with a library of 171 variant peptides where each residue (rows) was substituted with each other possible amino acid (columns) for 4 hours in serum-free media. (A) Cells were stained with PE-conjugated anti-HLA-E monoclonal antibody and surface levels of HLA-E were assessed through flow cytometry. (B) PBMCs were stimulated with anti-human CD3 monoclonal antibody and CD8-positive T cells were isolated and cultured without activating stimuli for 7 days. Pulsed cells from (A) were co-cultured with $CD8^{+}$ T cells and RLP-13. The concentration of chemokine MIP-1β in the co-culture supernatant was quantified through ELISA. The framed cells in the heatmaps correspond do the amino acid sequence in the original Mtb44 peptide.

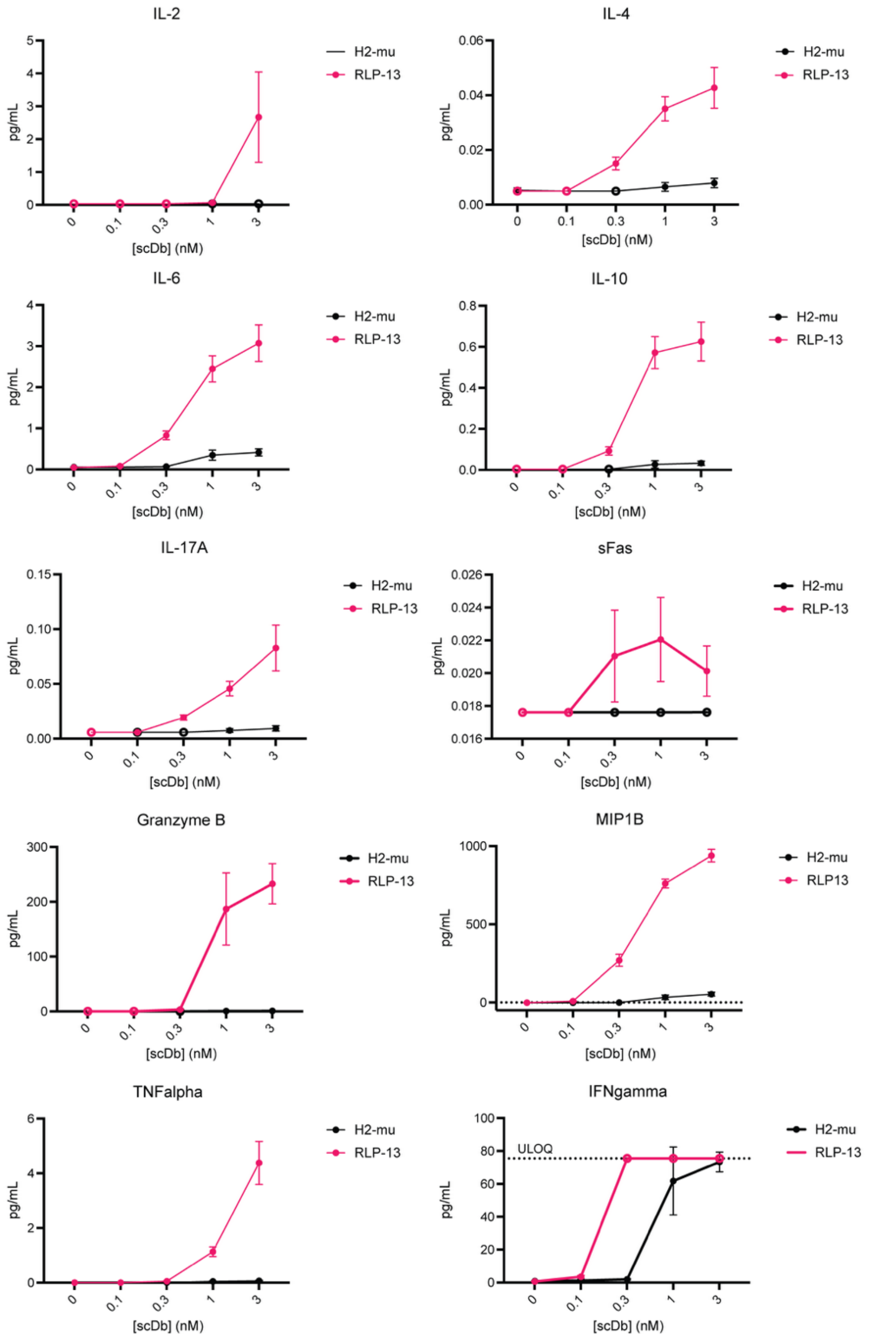


Supplementary Figure 5: Quantification of additional secreted cytokines, chemokines and effector molecules in co-culture supernatant from Figure 4C. Quantification performed using the BioLegend LEGENDPlex™ Human CD8/NK Panel (13-plex) kit following manufacturer protocol. ULOQ, upper limit of quantification. Black symbols: supernatant from co-culture with irrelevant control scDb; red symbols: co-culture with RLP-13. Empty symbols: values outside of quantifiable range (either below limit of detection or above ULOQ).

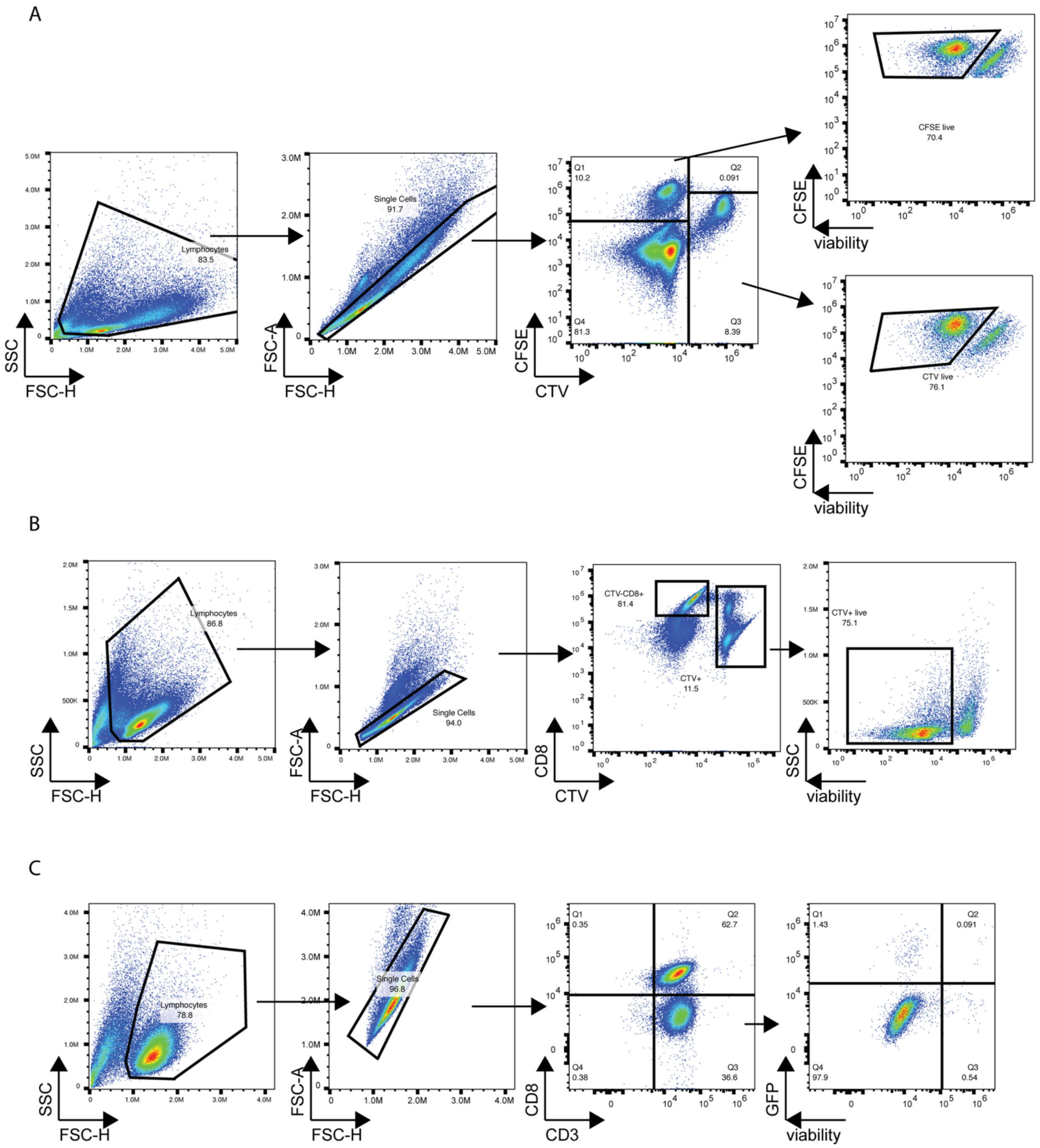


Supplementary Figure 6: Gating strategies for flow cytometry analysis of co-cultures. (A) Gating strategy for co-cultures in Figure 4C. (B) Gating strategy for co-cultures in Figure 5B. (C) Gating strategy for co-cultures in Figure 6D.

| Round | Phage Input Volume | Phage Output Volume | % Phage Used | Neg. Selection | | | Pos. Selection | | Notes |
|---|---|---|---|---|---|---|---|---|---|
| | | | | Bead Volume (uL) | pMHC Monomer | Cells (pulsed with indicated peptide) | Bead Volume (uL) | Mtb44-monomer (ug) | |
| 0 | 18mL | 18mL | 100 | 1000 | - | - | - | 0 | Neg. selected library against SA-beads |
| 1 | 5mL | 1 | 27 | 1000 | - | - | 50 | 1 | Pos. selection done twice, first with BAE wash, then with more stringent TBS-T (0.5%) wash |
| 2 | 100uL | 1 | 10 | 100 | 2ug (SP-2A, SP-1C, SQAPLPCVL, HIV.Gag275, SIV.Gag276) | 20M (SP-2A pulsed) and activated CD4T cells | 50 | 0.1 | |
| 3 | 10uL | 1 | 1 | 50 | 2ug (SP-2A, SQA-PLPCVL, SIV.Gag276, HIV.Gag275, SP-1A, EKQRESKREK, RIRTWKSLV, SALSEGATP, AISPRTLNA) | 120 M (SP-2A, RLPVKAPLL, RPLAAAPLL, RLPAKAPAL, activated CD4, activated Jurkat) | 30 | 0.01 | |
| 4 | 1uL | 1 | 0.1 | 50 | 1ug (SP-2A, SQA-PLPCVL, HIV.Gag275, SIV.Gag276, SP-1A, EKQRESKREK, RIRTWKSLV, SALSEGATP, AISPRTLNA) | 104M (SP-2A, ALPAKAPLL, RAPAKAPLL, RLAAKAPLL, and 7 M activated CD4) | 25 | 0.001 | |

Supplementary Table 1: Phage panning conditions for Mtb44/HLA-E*01:03 using the previously published Ludwig3 phage library.

| Round | Phage Input Volume | Phage Output Volume | % Phage Used | Neg. Selection | | | Pos. Selection | | Notes |
|---|---|---|---|---|---|---|---|---|---|
| | | | | Bead Volume (uL) | pMHC Monomer | Cells (pulsed with indicated peptide) | Bead Volume (uL) | HIV.-Gag275-monomer (ug) | |
| 0 | 18mL | 18mL | 100 | 1000 | - | - | - | 0 | Neg. selected library against SA-beads |
| 1 | 5mL | 1 | 27 | 1000 | - | - | 50 | 1 | Pos. selection done twice, first with BAE wash, then with more stringent TBS-T (0.5%) wash |
| 2 | 100uL | 1 | 10 | 100 | 2ug (SP-2A, SQA-PLPCVL, SIV.Gag276) | 20M (SP-2A pulsed) and activated CD4T cells | 50 | 0.1 | |
| 3 | 10uL | 1 | 1 | 50 | 2ug (SP-2A, SQA-PLPCVL, SIV.Gag276, SP-1A, EKQRESKREK, RIRTWKSLV, SALSEGATP, AISPRTLNA) | 120 M (SP-2A, SIV.Gag276, activated CD4, activated Jurkat) | 30 | 0.01 | |
| 4 | 1uL | 1 | 0.1 | 50 | 1ug (SP-2A, SQA-PLPCVL, SIV.Gag276, SP-1A, EKQRESKREK, RIRTWKSLV, SALSEGATP, AISPRTLNA) | 104M (SP-2A, SIV.Gag276, and 7M activated CD4) | 25 | 0.001 | |

Supplementary Table 2: Phage panning conditions for HIV.Gag275/HLA-E*01:03 using the previously published Ludwig3 phage library.

| Round | Phage Input Volume | Phage Output Volume | % Phage Used | Neg. Selection | | | Pos. Selection | | Notes |
|---|---|---|---|---|---|---|---|---|---|
| | | | | Bead Volume (uL) | pMHC Monomer | Cells (pulsed with indicated peptide) | Bead Volume (uL) | SIV.-Gag276-monomer (ug) | |
| 0 | 18mL | 18mL | 100 | 1000 | - | - | - | 0 | Neg. selected library against SA-beads |
| 1 | 5mL | 1 | 27 | 1000 | - | - | 50 | 1 | Pos. selection done twice, first with BAE wash, then with more stringent TBS-T (0.5%) wash |
| 2 | 100uL | 1 | 10 | 100 | 2ug (SP-2A, SQA, HIV.Gag275) | 20M (SP-2A pulsed) and activated CD4T cells | 50 | 0.1 | |
| 3 | 10uL | 1 | 1 | 50 | 2ug (SP-2A, SQA-PLPCVL, HIV.Gag275, SP-1A, EKQRESKREK, RIRTWKSLV, SALSEGATP, AISPRTLNA) | 120 M (SP-2A, HIV.Gag275, activated CD4, activated Jurkat) | 30 | 0.01 | |
| 4 | 1uL | 1 | 0.1 | 50 | 1ug (SP-2A, SQA-PLPCVL, HIV.Gag275, SP-1A, EKQRESKREK, RIRTWKSLV, SALSEGATP, AISPRTLNA) | 104M (SP-2A, HIV.Gag275, and 7M activated CD4) | 25 | 0.001 | |

Supplementary Table 3: Phage panning conditions for SIV.Gag276/HLA-E*01:03 using the previously published Ludwig3 phage library.
